# Distal super-enhancer drives aberrant *CXCL13* expression in Cancer cells driving growth and p53 dysregulation via CXCR5-CXCL13 axis

**DOI:** 10.1101/2024.08.31.609994

**Authors:** Santosh Kumar Gothwal, Pieta K. Mattila, Jacqueline H Barlow

**Affiliations:** Institute of Frontier Science Initiative, Kanazawa University, Japan; Central Research Resource Center, Cancer Research Institute, Kanazawa University; Institute of Biomedicine and MediCity Research Laboratories, University of Turku, Turku, Finland; InFLAMES Research Flagship, University of Turku, Turku, Finland; Turku Bioscience, University of Turku and Åbo Akademi University, Turku, Finland; Department of Microbiology and Molecular Genetics, University of California, Davis, United States

**Keywords:** CXCL13, super-enhancer, CXCR5, B-lymphoma, p53, GC-migration

## Abstract

The CXCL13 chemokine plays a crucial role in guiding B cell migration to the light zones (LZs) during the germinal center (GC) reaction. While *CXCL13* expression is absent in most cell types, aberrant amplification of the CXCR5-CXCL13 signaling is observed in various cancers, including germinal center-derived B-lymphomas (GCDBL), colorectal adenocarcinoma (COAD), and liver hepatocellular carcinoma (LIHC). However, the molecular mechanisms underlying abnormal *CXCL13* transcription in cancer cells and its functional consequences remain elusive. We identify DNA-CpG methylation binding protein 1 (MBD1) as a suppressor of *CXCL13* expression. Chromosomal conformation capture (3C) analysis reveals a distal super-enhancer located near *CCNG2* that interacts with the *CXCL13* promoter in GCDBL, suggesting that enhancer-hijacking drives the aberrant expression. Our functional validation demonstrates that CXCR5-CXCL13 signaling suppresses p53 and its target genes in GCDBLs, COAD, and LIHC. Notably, CXCL13 in the GCDBL cell line Raji disrupts CXCR5-mediated migration, a mechanism essential for (light zone) LZ-entry and affinity maturation of GC B cells. These findings highlight the dual role of the CXCR5-CXCL13 axis in immune response and cancer proliferation.

**Key Points:**

1. Super-enhancer near *CCNG2* region interacts with *CXCL13-TSS* driving CXCL13 in cancers.
2. Aberrant CXCL13 prevents CXCR5-mediated migration of B-lymphomas and promotes growth and p53 dysregulation in CXCR5+ cells
3. CXCR5-CXCL13 axis impairs p53 target gene expression and promotes tumor growth

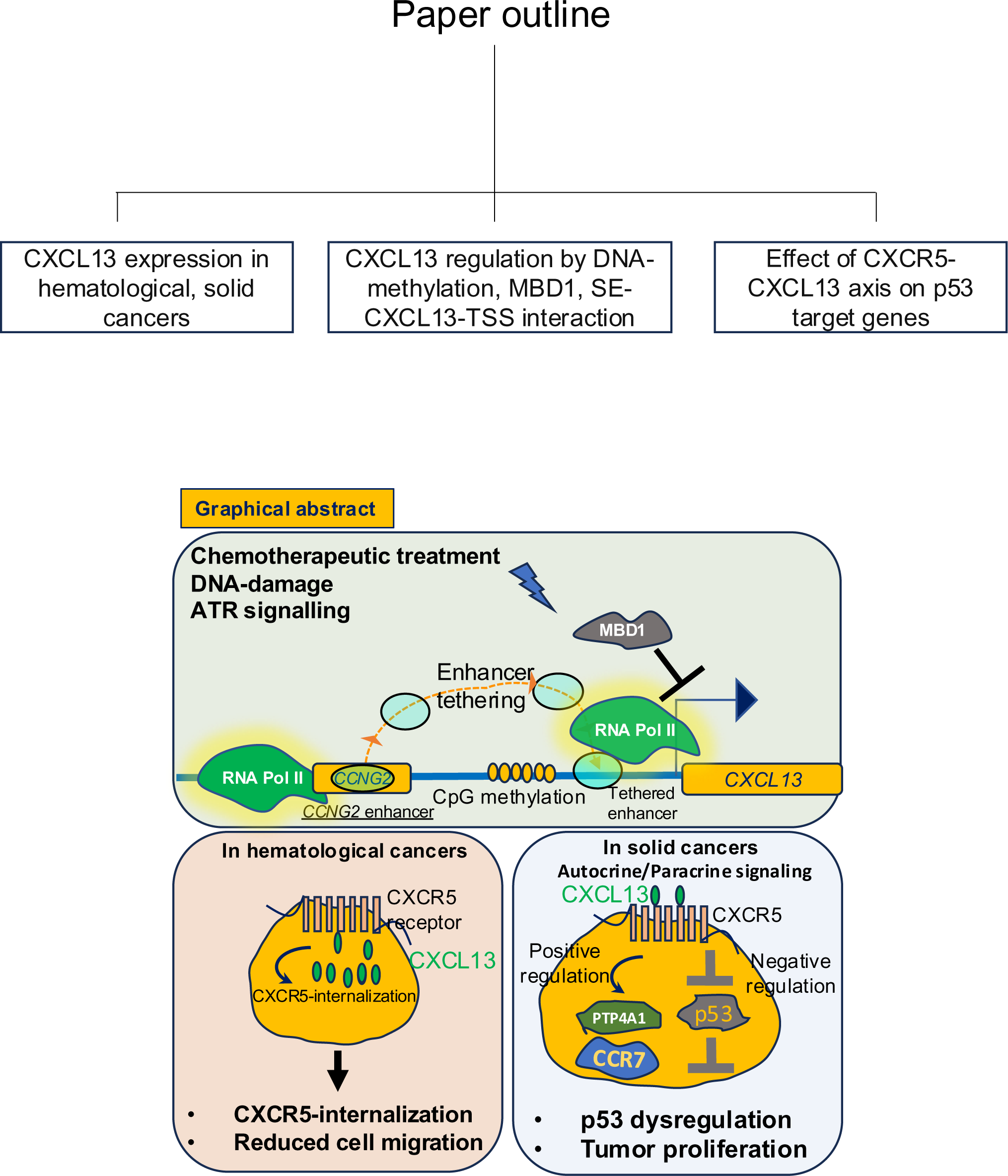

**Highlights:**

- Aberrant *CXCL13* expression in hematological and solid cancers
- Chemotherapeutic treatment of cancer cells promotes *CXCL13* and *CXCR5* expression
- Distal super-enhancer on *CCNG2* interacts with *CXCL13* promoter
- CXCL13 expression in B-lymphomas prevents CXCR5-dependent migration
- CXCR5-CXCL13 axis encounters p53 function in hematological and solid cancer cells

## Introduction

Chemokine (C-X-C motif) ligand 13 (CXCL13) binds and exerts signaling through the C-X-C chemokine receptor type 5 (CXCR5) and G proteins [1–4]. *CXCL13* is primarily expressed in follicular dendritic cells (FDCs), follicular helper T cells (Tfh), CD4+ T cells, CD8+ T cells, stromal cells and macrophages and regulates the innate and humoral immune repsonse [1–3, 5–9]. But aberrant *CXCL13* expression is also observed in hematological malignances such as B-cell chronic lymphocytic leukemia (B-CLL) and multiple myeloma (MM) [10–13]. Among solid cancers, *CXCL13* expression is observed in non-small cell lung carcinoma (NSCLC), breast cancers, prostate cancers and *Helicobacter pylori* associated gastric lymphomas [11, 14–18]. Aberrant CXCR5-CXCL13 signaling is evident in autoimmune diseases such as rheumatoid arthritis, [19, 20], multiple sclerosis [21], systematic lupus erythematosis [22], primary Sjögren’s syndrome [23] and myasthenia gravis [24, 25].

CXCL13 signaling is known to interfere the chemotherapeutic response and elevated CXCL13 expression by cancer cells promotes autocrine and paracrine effects in the tumor microenvironment resulting [26]. CXCL13 secreted by MM cells further induces CXCL13 in surrounding macrophages by Bruton’s tyrosine kinase (BTK) signaling. In turn, macrophages induce *CXCL13* expression in MM through TGF-β signaling [13]. Benzo(a)pyrene (BaP), a carcinogen found in tobacco and air pollution, induces *CXCL13* in NSCLC [27]. *CXCL13* knockdown in MM and NSCLC resulted in slower tumor growth in xenograft experiments [13, 27]. The CXCR5-CXCL13 axis exhibits resistance to chemotherapeutic drugs including bortezomib in MM [28], and 5-flurouracil in colorectal cancer [29] and mantle cell lymphoma [30]. In breast cancer, CXCR5-CXCL13 axis promotes metastasis by regulating the epithelial to mesenchymal transition (EMT) [31]. CXCR5-CXCL13 signaling promotes tumorigenesis in *PTEN* deficient cancers by Protein kinase C (PKC) signaling [32]. Loss of *PTEN* can induce CXCL13 expression by NF-kB signaling [32]*. PTEN* is a p53 dependent gene, suggesting that CXCR5-CXCL13 signaling interferes with p53 regulation, however whether the CXCR5-CXCL13 axis promotes tumorigenesis by interfering with other p53 target genes is not known.

Despite the widespread expression of *CXCL13* in hematological and solid cancers, the mechanism(s) driving aberrant *CXCL13* expression in cancer cells is elusive. Little is known about the factors directly regulating the transcriptional axis of *CXCL13*. TGF-β-induced SOX4 promotes the *CXCL13* expression during Th2 cells differentiation and retinoic acid and neuronal signaling induces the CXCL13 expression in murine embryonic stromal cells [7, 33]. Enhancer tethering has been suggested as a key mechanism for oncogene expression in cancers {Bi, 2020 #127;Dixon, 2012 #121;Roe, 2017 #128. Whether the CXCL13 promoter interacts with a distal enhancer that drives *CXCL13* expression in cancer cells remains unknown. The chromatin insulating protein CTCF links chromosome conformation and global gene-expression [34]. CTCF binding to DNA is driven by local DNA methylation levels and Ten eleven translocation (TET) enzymes [35, 36]. DNA CpG methylation and MBD1 drive transcriptional suppression [37, 38]. Whether MBD1, TET enzymes, and CTCF regulate *CXCL13* expression remains unclear.

In this study, we identify a super-enhancer, surrounding the *CXCL13* locus interacting with *CXCL13* locus in cancer cells. Our findings in the human Burkitt’s lymphoma cell line Raji, suggest a role for aberrantly expressed CXCL13 in preventing migration of GCDBL cells towards external CXCL13, a LZ chemokine. CXCR5-CXCL13 axis suppressed p53 target genes in B-lymphomas, COAD and LIHC. These findings highlight the precise control of *CXCL13* expression through epigenetic mechanisms and the regulation of enhancer-promoter interactions.

A loss of these regulatory processes in cancer is associated with the complex roles of CXCR5-CXCL13 signaling in both hematological and solid tumors. Our study highlights the importance of CXCL13 regulation in humoral immunity and tumorigenesis.

## Material and Methods

### Human B lymphoma cultures and treatments

Human B lymphoma Cell lines FL-18, FL-218, FL-318, FL-518, Raji and Granta-519 were obtained from RIKEN, Japan and Kyoto university hospital. All B lymphoma cell lines, THP1, LoVo and HC116 were cultured in RPMI (30264-85, Nakalai) containing 10% FBS, 1% Penicillin, Streptomycin and Amphotericin B solution (Wako# 161-23181) in CO_2_ incubator at 37°C under 5% CO_2_ concentration. MCF7, HepG2 and 293T cells were cultured in the DMEM containing 10% FBS, 1% Penicillin, Streptomycin and Amphotericin B solution (Wako# 161-23181) in CO_2_ incubator at 37°C under 5% CO_2_ concentration. Treatment of cells with chemotherapeutics drugs Hydroxyurea (Product #085-06653, Wako, Japan), Gemcitabine (product # S1714, Selleck, Japan) ATR inhibitor (catalogue#KU55933; Selleck, Japan), NF-κB inhibitor (catalogue## S4902: Selleck, Japan), AKT inhibitor (catalogue#S1037:Selleck, Japan) and etoposide (product #055-0843, Wako Japan), bobocat339 (catalogue #SML2611, sigma), Bleomycin (Catalogue# B7116-1ML, Sigma), Nutlin-3a (Catalogue# S8059, Selleck, Japan) were used as per defined timepoints and concentrations.

### Short hairpin knockdown, and lentivirus preparation

293T cells at nearly 60% confluency were transfected with a mix of *p-PAX2* (3 µg), *pVSVG* (3 µg), and target shRNA plasmids in *pLKO1-puromycin/blasticidin backbone* (3 µg). Plasmids were mixed in 1 ml of Opti-Mem (Cat# 31985062, Gibco), then 27 µl of Polyethyleneimine (PEI) solution (1 µg/ml) was added. After 20 minutes at room temperature, the mixture was used for transfecting 293T cells. Cells were refreshed with DMEM medium after 24 hours, and culture supernatants were collected 48 hours post-transfection, filtered with a 0.2 µM filter. Lentiviral medium (1 ml) was used for transduction to 1 million Raji cells, followed by puromycin (1 µg/ml) or Blasticidin (10 µg/ml) selection for resistance. Puromycin or Blasticidin was refreshed every three days for one week. For production of CRISPR/Cas9 lentivirus, guideRNA cloned into the Guide-it™ CRISPR/Cas9 Systems (Takarabio# 632601) were used instead of shRNA plasmids follwed by the similar procedure and transduction to Raji cells.

### B Cell Migration Assay

Transwell (Costar, REF#3421) was filled with RPMI (5% FBS) or RPMI (5% FBS) medium containing rhCXCL12 (200 ng/ml) or rhCXCL13 (1000 ng/ml) (Biolegend). Cell suspension of 5*10^5^ cells or 1.5*10^5^ cells for normal human B cells, isolated by institutional approval and with manufacturers instruction, EasySep™ Human B Cell Isolation Kit (STEMCELL technologies) were placed in transwell inserts for 6 hours at 37°C in CO_2_ chamber. Collected cells were lysed in 200 μl of CellTiter-Glo® 2.0 Reagent (Promega #G9242) and allowed to stabilize for 10 minutes to measure luminescence signals. Absorbance was measured using TECAN luminometer (code # MIC9414). A CXCL13-migration rescue experiment was conducted in Raji cells, transfected with lentiviral particles containing CRISPR/Cas9 guide RNA targeting exons 2 and 3 of *CXCL13*. Following transfection, cells were cultured for 4 days before being utilized in a migration assay.

### FACS analysis

Raji cells (1 * 10^6) were surface-stained with isotype controls or antibodies (Supplementary Table 1) constituted in FACS buffer (PBS from Nakalai Tasque, 0.1% BSA, 0.1% NaN3). After a 15-minute staining at room temperature, cells were washed, resuspended in 400 μl FACS buffer, and mixed with 7-AAD before analysis. Flow cytometry was conducted using a FACS Fortessa Flow cytometer (Becton Dickinson, San Jose, USA), analyzing 10,000 events with data displayed as two-color plots (SSC-A/APC/PE/BV-421). Regarding CRISPR/Cas9 editing, Raji cells were transfected with guide RNA targeting *CXCL13* exons 2 and 3, then sorted using a ZSgreen marker.

### Cloning, sequence analysis and overexpression

Full-length CXCR5 transcript amplification was performed from Raji’s cDNA with primers listed in supplementary Table 1. The PCR product was digested with NheI and BamHI and cloned *pCMV-IRES-Puro vectors*, Sequenced and, aligned with Uniprot accession (P32302). 5 µg of HpaI-linearized *pCMV-CXCR5-IRES-Puro* was transfected into 1 million Raji cells via Nepagene 21 electroporation followed by Puromycin selection (1 µg/ml) for a week to obtain stably transfected cells. Stable integration was confirmed by PCR, flow cytometry and qPCR. The Nepagene 21 electroporation system was used according to the instructions for the transient transfection of *pcDNA-MBD1*, *pcDNA-BCL6*, and *pcDNA* into cell lines.

### RNA isolation, cDNA preparation, RT-qPCR analysis and RNA-seq datasets from GEOR

RNA from of normal Human B and B lymphoma cells was isolated (NucleoSpin® RNA kit, #740984.50, Takara), cDNA was prepared (FSQ#201, Toyobo) and RT-qPCR was performed with SYBR qPCR mix (QPS-201, Toyobo). The primer sequence of the analyzed transcripts is provided (Supplementary table 1). Standard delta-delta Ct method was used for analysis of relative transcript levels as described [39]. GSE171763 and GSE190319 datasets were used from GEO2R, NCBI. Accession token (gfkvukuqnpsfvmx) and accession number GSE236240 were utilized fot the cMYC and p53 ChIP-seq peaks in Raji.

### Immunoprecipitation and Immunoblotting

2 million Raji cells were collected, washed with PBS and lysed in 200 μl of RIPA buffer containing 1X protease inhibitors cocktail (Sigma#11873580001). Lysed cells were spun at 13,000g for 10 min at 4 °C. The cell lysate from supernatant was used for IP with 3 μg of CXCL13/IgG antibody (AF801, R&D system) and 20μl precleared protein A beads (Invitrogen; 10001D, Invitrogen) for 2 hours at room temperatures, followed by washing of the beads with RIPA buffer up to three times, 5 minutes each. The washed beads were used for the elution and protein blotting. For Western blotting, cleared cell lysates from 1 million cells (100 μl of RIPA) was reconstituted in 1X Lamellae buffer (Bio-Rad) with a final concentration of 10% β-mercaptoethanol, followed by boiling at 95 °C for 5 min. After SDS SDS-PAGE and transferring to a PVDF membrane using the Transblot® SD Semi-Dry Electrophoretic Transfer Cell (Product #1703940, Bio-Rad), PVDF membranes were blocked in 5% skim milk for 15 min, washed with 1X TBST, and probed with primary antibody solution overnight at 4 °C. The following day, the membrane was washed three times with 1X-TBST (10 min each) and then incubated with an HRP-conjugated secondary antibody for 1 hr. at room temperature. After washing, the membrane was used to develop a chemiluminescent signal using the Chemi-lumi one ultra (product #11644-40, Nakalai, Japan). The protein qualification was performed using the ImageJ.

### Chromosome Conformation Capture (3C) Assay

The 3C assay was performed as described [39]. Two million Raji cells (-/+HU) were fixed in 1% PFA/PBS for 10 minutes, quenched with glycine, and lysed in 500μl buffer (20 mM Tris-HCl, pH-8; 0.5% NP40; 1 mM EDTA, pH-8, 1X protease inhibitor, Sigma #11873580001) for 30 minutes. 760 μl of 1X CutSmart buffer (NEB #B6004) and 24μl of 10% SDS buffer were added, incubated at 37°C for 1 hour, followed by EcoRI-digestion (40μl), overnight. The next day, 80 μl of 10% SDS was added, incubated at 65°C (20 minutes), then mixed with 9.2 ml of 1X ligation-buffer, 100 μl of Triton X-100, and 1750 units of Hi-T4™ DNA Ligase (NEB#M2622S). Ligation was performed at 16°C for 5 hours and addition 16-18 hours at room-temperature. Samples were treated with 10 uL of RNase A (10 mg/ml) and 20 uL of proteinase K (20 mg/ml) before DNA purification. The DNA pellet was dissolved in 600 μl TE buffer, 1μg of DNA was used for PCR (Takara, HS-Taq#R007A).

### Chromatin Immunoprecipitation (ChIP)

ChIP assay was performed as described earlier [39]. Briefly, 2 × 10^7^ cells were crosslinked in 50 ml of PBS with 1% formaldehyde at room temperature, quenching with 2 ml of 2M glycine, followed by PBS washing, cells lysis (20mM Tris-HCL (pH-8), 0.5% NP40, 1mM EDTA (pH-8), with 1X protease inhibitors, NaF (20 mM), and Na3Vo4 (1 μM). After brief centrifugation and nuclei collection, they were treated with SDS and 8M urea buffers, followed by sonication (13 cycles, 30 sec on, 1 min off, Diagenode). Approximately 25 μg of chromatin was immunoprecipitated with 2 μg of antibodies, and 25 ul of Protein A/G Magna ChIP beads (Sigma). The eluted samples and Input DNA were treated with RNAse and proteinase K, then purified for qPCR analysis of target promoters. Primers and antibodies for qPCR are shown in (supplementary table 1 and 2).

## Results

### Aberrant *CXCL13* expression in hematological malignancies and solid cancers

To identify cancer types exhibiting aberrant *CXCL13* expression, we explored the cancer genome atlas (TCGA) sorting tumor samples by altered *CXCL13* status including deep deletion, shallow deletion, diploid, gain, and amplification [40–42]. Genomic Identification of Significant Targets in Cancer (GISTIC) analysis of *CXCL13* copy number alterations confirmed higher log2 copy number values in gain and amplified groups compared to deep deletion, shallow deletion, and diploid groups (Figure 1A). However, the levels of *CXCL13* mRNA in *CXCL13*-gain, *CXCL13*-amplified, *CXCL13*-deletion and *CXCL13* shallow-deletion groups were not significantly different from *CXCL13*-diploid control groups (Figure 1B). This suggests that genetic alterations such as amplification and gains leading to increased copy number of *CXCL13* are not essential determinants of enhanced *CXCL13* mRNA expression in cancers (Fig 1A-B) and implies that the normally tight regulation of *CXCL13* expression becomes leaky in specific cancers.

**Figure 1.**
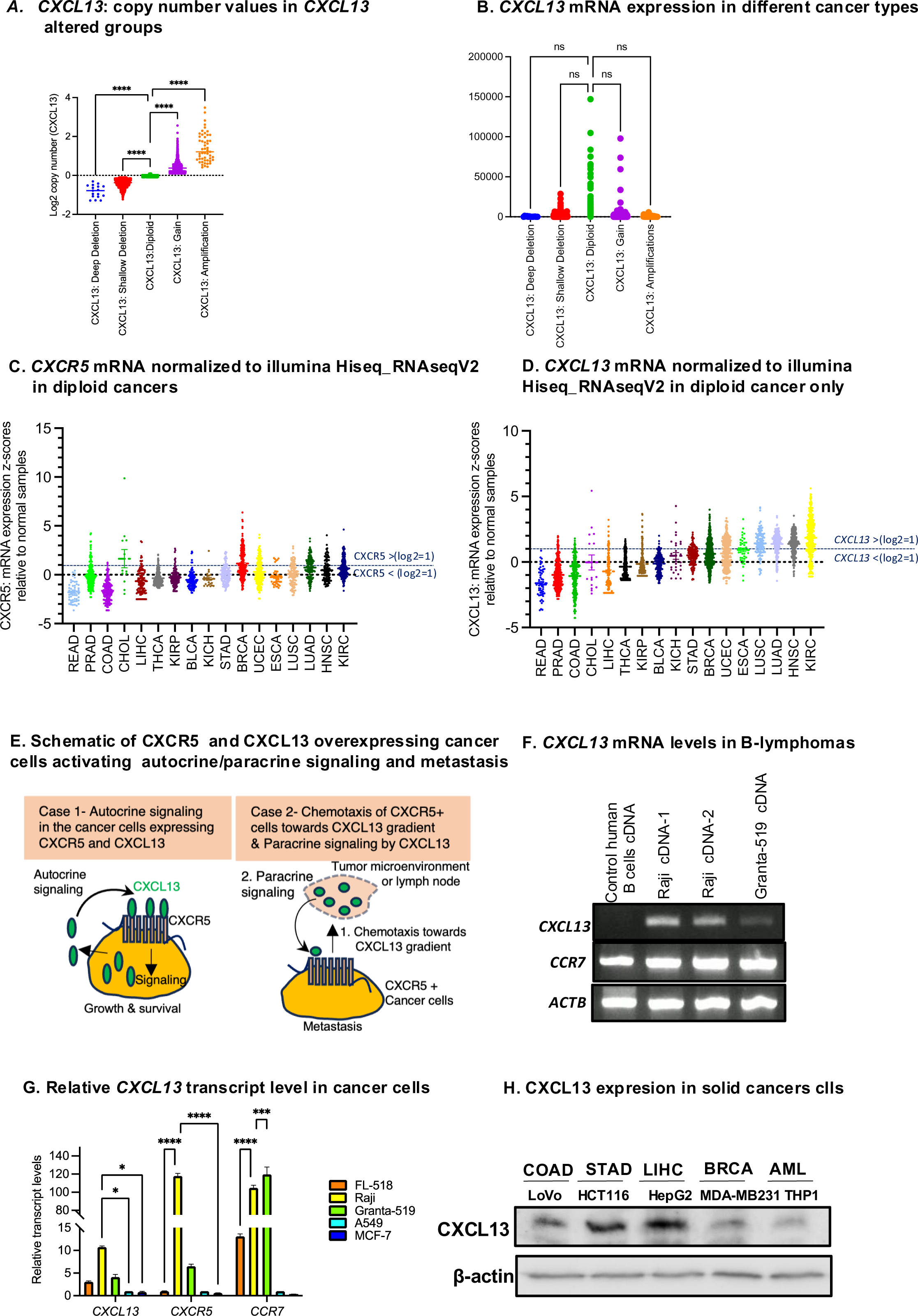
*CXCL13* is expressed in hematological, and solid cancers. (A) *CXCL13* putative copy number values from GISTIC vs log2 copy number values in 10712 samples. p values <0.0001 are derived from ordinary one-way ANOVA (B) *CXCL13*: mRNA expression z-scores relative to normal samples (log RNA-Seq V2 RSEM) data are from 6909 samples in both axes. *ns* = not significant; p value derived from ordinary one-way ANOVA (C) *CXCR5*: mRNA expression z-scores relative to normal samples (log2 RNA Seq V2 RSEM from 6959 diploid samples *RSEM= RNA-seq* by expectation-maximization. X axis profile shows TCGA cancer acronyms. Dotted line in parallel to X axis separates the tumor samples exhibiting cutoff *CXCR5 mRNA* log2 values >1 or log2<1. TCGA Cancer acronyms: rectum adenocarcinoma (READ), bladder urothelial Carcinoma (BLCA), breast cancer (BRCA), cholangiocarcinoma (CHOL), colon adenocarcinoma (COAD), esophageal carcinoma (ESCA), head and neck squamous cell carcinoma (HNSC), kidney chromophobe (KICH), Kidney renal clear cell carcinoma (KIRC), kidney renal papillary cell carcinoma (KIRP), liver hepatocellular carcinomas (LIHC), lung adenocarcinoma (LUAD), lung squamous cell carcinoma (LUSC), prostate adenocarcinoma (PRAD), stomach adenocarcinoma (STAD), thyroid carcinoma (THCA), uterine corpus endometrial carcinoma (UCEC) (D) *CXCL13*: mRNA expression z-scores relative to normal samples (log RNA Seq V2 RSEM) in 6953 diploid samples. X axis shows TCGA cancer acronyms. Dotted line in parallel to X axis separates the tumor samples exhibiting cutoff *CXCL13 mRNA* log2 values >1 or log2<1 (E) Schematic of CXCR5 and CXCL13 overexpressing cancer cells in activating autocrine, paracrine signaling, and metastasis to distally located CXCL13 chemokine gradient. (F) Ethidium bromide gel-electrophoresis of *CXCL13* PCR using the cDNA samples from normal human peripheral B cells, Raji and Granta-519 cells (G) Relative quantitative real-time PCR (qRT-PCR) analysis comparing the mRNA levels of *CXCL13* versus *ACTB* in Raji, FL-518, Granta-519, A549, and MCF7 cells. Additionally, quantification of *CXCL13*, *CXCR5*, and *CCR7* mRNA levels in multiple human B cell lymphoma cell lines (Raji, FL-518, Granta-519), along with breast cancer (MCF7) and lung cancer (A549) cell lines. *CXCL13* mRNA levels in the other cell lines are normalized to those in A549 cells. Turkey’s multiple comparison test for statistical significance. *p= 0.0486* for Raji cells vs A549 cells and *p= 0.0430* for Raji cells vs MCF-7 cells. Data are mean ± SEM for n=3 (H) CXCL13 Western blotting using LoVo (COAD), HCT116 (COAD), HepG2 (LIHC), MDAMB-231 (BRCA) and THP1 (Acute monocytic leukemia) cell lysates.

To examine the expression of *CXCL13* mRNA among various cancers, we plotted *CXCR5* and *CXCL13* mRNA Z-scores normalized with normal samples among cancer types (Figure 1C-D). For both *CXCR5* and CXC13, mRNA log2 RNA-seq vs RSEM values above log2 cutoff value of 1 were found in at least 1 sample for all cancers analyzed except for rectum adenocarcinoma (READ) (Figure 1C,D). These results suggest that both the CXCR5 receptor and CXCL13 ligand are potentially co-expressed in several cancers and may promote autocrine signaling (Figure 1C-E). To evaluate whether *CXCL13* and *CXCR5* are upregulated in the same tumor, we randomly identified seven CHOL samples exhibiting *CXCR5* mRNA log2 value of more than 0 (Supplementary Figure 1A). Four out of seven samples also exhibited *CXCL13* expression (Supplementary Figure 1A), demonstrating that CHOL exhibits co-expression of *CXCR5* and *CXCL13* and suggests co-upregulation may occur in multiple cancer types (Figure 1C-E, supplementary Figure 1A). Downstream signaling by the CXCR5-CXCL13 axis in these cancer types may promote therapy resistance (Figure 1C-E). Of note, the overall survival (OS) was notably lower among high-CXCL13 expressors compared to those with lower *CXCL13* levels as seen across Acute Myeloid Leukemia, Glioblastoma, and Uveal Melanoma (Supplementary Figure 1B-D).

Based on widespread *CXCL13* expression in human cancers (Figure 1A-D), we examined *CXCL13* expression in human cell lines derived from hematological and solid malignancies. *CXCL13* mRNA and protein expression was confirmed among Raji (Burkitt’s lymphoma), Granta-519 (mantle cell lymphoma), FL-518 (Follicular Lymphoma), A549, MCF7, HepG2 (LIHC), HCT116, and LoVo (COADs) MDA-MB231 and THP1 cells (Figure 1F-H). These observations support the notion of aberrant *CXCL13* expression in hematological and solid cancers.

### Induced *CXCL13* expression after chemotherapeutic treatment of cancer cells targeting DNA replication, DNA-damage and BTK signaling

Chemotherapeutic treatments in MM and NSCLC induce *CXCL13* and therapy resistance [13, 27]. To determine whether chemotherapeutics that induce DNA damage response and inhibit DNA replication affect *CXCL13* expression in cancer cells, we investigated the effects of hydroxyurea (HU), gemcitabine, etoposide, and bleomycin on *CXCL13* expression (Figure 2A-D). In Raji and HCT116 cells, exposure to HU increased CXCL13 expression (Figure 2A, supplementary figure 2A). In contrast, γH2AX, a marker of DNA damage, was decreased in Raji cells treated with 20 mM HU than cells treated with 10 mM HU (Figure 2A, B). Similarly, treatment of Raji cells with gemcitabine, etoposide and bleomycin increased the CXCL13 levels however no reduction in γH2AX was observed upon this treatment (Figure 2A-D). Taken together, these results show that DNA damage induced by various chemotherapeutics induces CXCL13 expression (Figure 2A-D). It is possible that higher CXCL13 levels may counteract DNA replication stress-induced DNA damage caused by HU treatment (Figure 2B). These suggest a possibility to check in future whether CXCR5-CXCL13 axis promotes the γH2AX turnover and DNA-repair.

**Figure 2.**
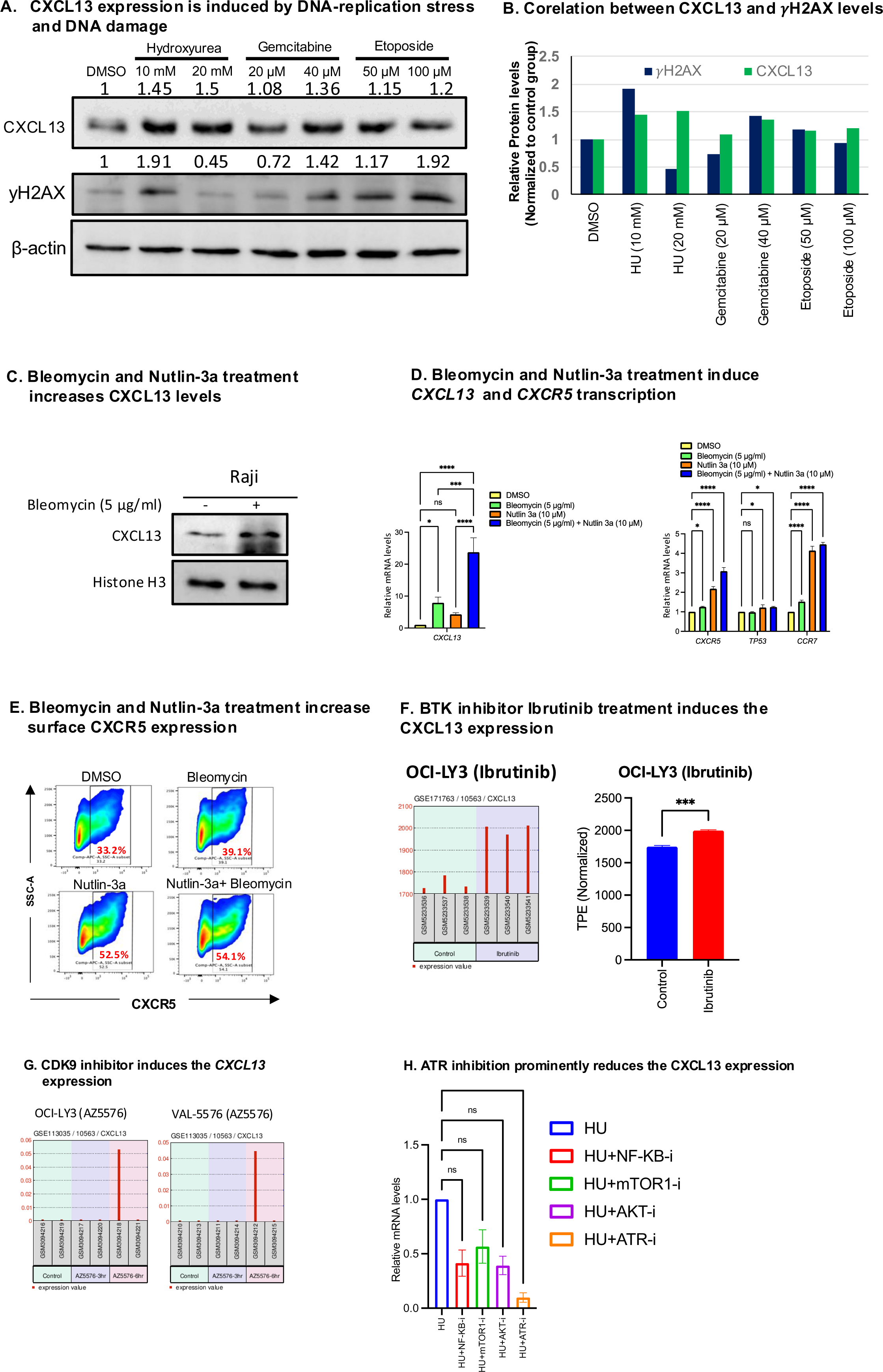
DNA damage and Replication stress induce *CXCL13* expression in the Raji cells. (A) Western blotting of CXCL13 and γH2AX from the cell lysates of Raji cells treated with DMSO, hydroxyurea (10 mM and 20 mM), Gemcitabine (20μM and 40μM) and Etoposide (50μM and 100μM) for 24 hours. Values above blot indicate densitometric qualification of CXCL13 normalized to β-actin levels from a representative blot (B) Correlation of densitometry quantification of CXCL13 and γH2AX signals in Raji cells treated with HU, Gemcitabine and Etoposide, normalized with CXCL13/γH2AX levels of DMSO treated Raji cells (C) CXCL13 expression analysis by Western blotting in Raji cells treated with Bleomycin (5μg/ml) for 24 hrs. Values above blot indicate densitometric qualification of CXCL13 normalized to β-actin levels from a representative blot (D) qRT-PCR analysis of *CXCL13*, *CXCR5*, *TP53* and CCR7 mRNA levels in Bleomycin (5μg/ml) treated Raji cells, Nutlin-3a (10μM) treated Raji cells and Bleomycin (5μg/ml) + Nutlin-3a (10μM) treated Raji cells are shown. ***CXCL13***: p= 0.0409 in Raji (DMSO vs Bleomycin), p= 0.5163 (DMSO vs Nutlin-3a), p= 0.0002 [bleomycin vs (Nutlin-3a+Bleomycin)], p <0.0001 for [Nutlin-3a vs (Nutlin-3a+Bleomycin)]. Šídák’s multiple comparisons test. ***CXCR5***; p <0.0001 [DMSO vs (Nutlin-3a+Bleomycin)]. ***CCR7***; p <0.0001 [DMSO vs (Nutlin-3a+Bleomycin)]. Dunnett’s multiple comparisons test. Data are presented as mean ± SEM; n=3; p<0.0001 for DMSO versus Bleomycin treated groups, p<0.0001 for DMSO versus Nutlin-3a (10μM), and p<0.0001 for DMSO vs co-treatment with Bleomycin (5μg/ml) and Nutlin-3a (10μM). (E) Surface expression of CXCR5 in Raji cells in Raji cells treated with Bleomycin (5μg/ml), Nutlin-3a (10μM), and the combination of Bleomycin (5μg/ml) + Nutlin-3a (10μM). (F) Treatment with Bruton Tyrosine Kinase Inhibitor Ibrutinib to OCI-LY3 for 48 hrs induced *CXCL13* transcripts per million reads (TPM). Unpaired t-test, *p = 0.0004* (G) Panel of OCI-LY3 and VAL cell lines treated with AZ-5576, a selective CDK9 inhibitor, exhibiting induced *CXCL13* TPM (GSE113035) in at least 1/2 independent samples. (H) effect of 24 hr treatment of NF-κB inhibitor (10mM), AKT inhibitor (10μM), ATR inhibitors (10μM), mTOR1C (10μM) treatment on *CXCL13* mRNA expression in the Raji cells. Data are mean ± SEM of three independent experiments. Ordinary one-way ANOVA. *p*= 0.0118 for *HU vs HU + ATRi* groups.

DNA damage and DNA replication-stress induced signaling pathways activate p53 [43]. Therefore, we tested if p53 activation promotes *CXCL13* expression (Figure 2D). We examined the effect of Nutlin-3a treatment, an agonist of p53, on *CXCL13* levels in Raji cells (Figure 2D). Nutlin-3a treatment induced the p53-dependent gene *MDM2*, but at a lower level than in A549 cells which harbors a wild type p53 (Supplementary Figure 2 A, B), suggesting that Nutlin-3a partially activates the mutant p53 present in Raji cells (Supplementary Figure 2A, B). Intriguingly, treatment with either Nutlin-3a or bleomycin further increased *CXCL13* levels as well as *CXCR5* and *CCR7*, the GPCRs expressed in GC B cells (Figure 2D, E). Surface expression of CXCR5 in Raji cells was induced upon bleomycin and Nutlin-3a treatment (Figure 2E). These results suggests that chemotherapeutic treatment also induce CXCR5 expression in cancer cells. These results also imply a role for mutant p53 in regulation of *CXCL13, CXCR5 and CCR7* expression (Figure 2D-E). In addition, treatment with Ibrutinib (a BTK inhibitor) and AZ5576 (a CDK9 inhibitor) led to an upregulation of *CXCL13* expression in OCI-LY3 cells, which are an activated germinal center B cell-like diffuse large B-cell lymphoma (AGC-DLBCL) cell line, as well as in VAL cell line (B-lymphoma) (Figure 2F, G) [44]. This suggests that chemotherapeutics agents inhibiting the BTK signaling and CDK9 inhibitors induce the *CXCL13* and CXCR5 expression in hematological and solid cancers. Cancer cell survival is typically promoted by increased NF-kB, mTOR1C, AKT and ataxia telangiectasia and Rad3-related protein (ATR) signaling [45]. To examine if these pathways affect *CXCL13* expression under DNA-replication stress, we measured *CXCL13* mRNA by qRT-PCR in Raji cells treated with inhibitors against NF-kB, mTOR1C, AKT, and ATR (Figure 2H) in the HU-treated Raji cells (Figure 2H). The profound reduction on *CXCL13* mRNA levels was seen in response to ATR inhibition (Figure 2H). Overall these suggests that chemotherapeutics agents inhibiting the DNA-replication, DNA-damage response, BTK signaling and CDK9 inhibitors induce the *CXCL13* and CXCR5 expression in hematological and solid cancers while ATR signaling promotes the CXCL13 signaling.

### MBD1 regulates the *CXCL13* transcription independently of CTCF

By *in silico* analysis of a eukaryotic promoter database, we found putative binding sites for CTCF, retinoic acid receptor alpha (RARA), p53, and MYC in the 1 kilobase upstream of the *CXCL13* transcriptional start site (TSS) (Figure 3A-D). These factors also exhibited binding on the mouse *CXCL13* promoter (Supplementary figure 3A-C), suggesting an evolutionary conserved regulation of *CXCL13* by these factors (Figure 3A-D, Supplementary Figure 3A-C). MBD1 physically interacts with RARA and promyelocytic leukemia protein (PML), leading to histone deacetylase-3 mediated gene silencing [46]. Given the association among MBD1, RARA and CTCF with DNA-methylation, we tested the effect of MBD1 and CTCF on *CXCL13* expression (Figure 3E-F). Short hairpin (shRNA) silencing of *MBD1* and *CTCF* was confirmed in Raji cells (Figure 3F). Interestingly, shMBD1-Raji cells induced *CXCL13* expression compared to scramble control (shSCR-Raji) (Figure 3E), while shCTCF-Raji cells exhibited slight reduction in *CXCL13* mRNA levels (Figure 3E). Compared to shMBD1 alone, shMBD1+shCTCF-Raji cells exhibited a higher reduction in *CXCL13* expression (Figure 3E). These results suggest a suppressive role of MBD1 on *CXCL13*-expresion but a positive role of CTCF in *CXCL13* expression in MBD1-depleted conditions (Figure 3E).

**Figure 3.**
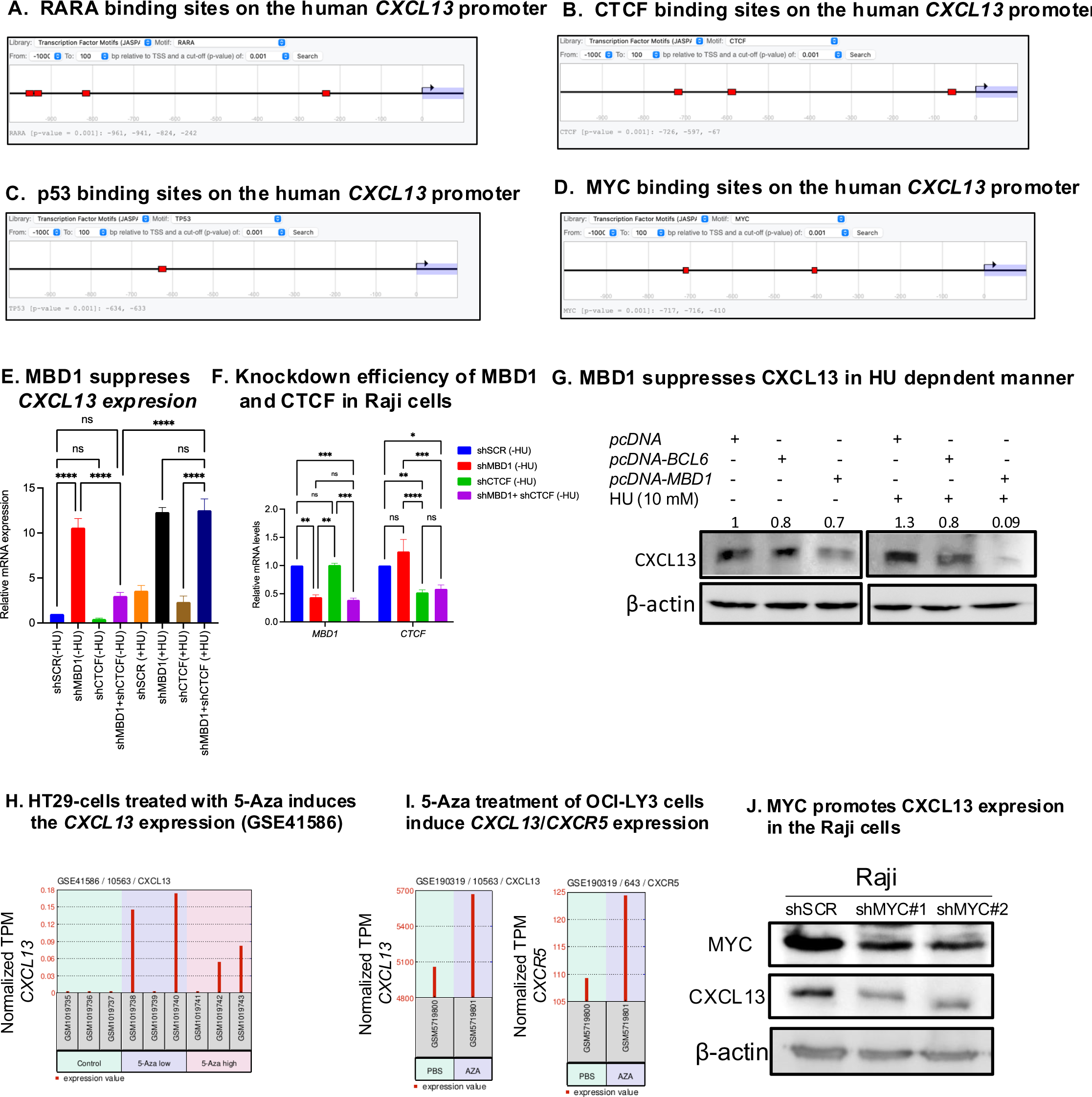
MBD1 suppresses the *CXCL13* expression independently of CTCF. (A) *In silico* binding of RARA, MYC, p53 and CTCF on *CXCL13* promoter predicted by eukaryotic promoter database. Factors binding to promoter with a cutoff p value > 0.001 are shown. Binding sites of RARA at −242, −824, −941, −961 from the *CXCL13*-TSS (B) Binding sites of CTCF at −67, −597, - 726 from TSS (C) Binding sites for p53 at −633, and −634 from TSS (D) Binding sites for MYC at −410, −716 and −717 from TSS (E) qRT-PCR quantification of *CXCL13* mRNA in -/+ HU treated shSCR-Raji, shMBD1-Raji, shCTCF-Raji and (shMBD1+shCTCF)-Raji cells. Data are mean ± SEM for three independent replicates. *p* values are derived from ordinary one-way ANOVA (*p<0.0001=****)* (F) qRT-PCR analysis of the *MBD1* and *CTCF* in shSCR-Raji, shMBD1-Raji, shCTCF-Raji and (shMBD1+shCTCF)-Raji cells. Data are mean ± SEM for three intendent replicates. *p* values are derived after Šídák multiple comparison test (G) CXCL13 Western blotting using Raji cell lysates transfected with *pcDNA-3.1*, *pcDNA-MBD1* and *pcDNA-BCL6* for 24 hours followed by 10 mM HU treatment for 24 hours. Cell lysates were prepared 48 hrs post-transfection and 24 hr. post HU treatment and then subjected to *CXCL13* western blotting. Values above blot indicate densitometric qualification of CXCL13 normalized to beta-actin levels from a representative blot (H) HT29-cells treated with 5-Aza induces the *CXCL13* expression. HT-29 cells treated with dimethyl sulfoxide (vehicle alone= 0 μM 5-Aza), 5μM 5-Aza and 10 μM 5-Aza for five days. Normalized transcripts per million (TPM) values of the *CXCL13* is shown. Two treated sample out of three samples exhibited the upregulation of *CXCL13* transcript (I) 5-Azacytidine treatment of OCI-LY3 cells induced *CXCL13* and *CXCR5* transcript levels (GSE190319). Normalized TPM values are shown in the graph (J) WB of CXCL13 using the cell lysates from shSCR-Raji, shMYC#1-Raji, shMYC#2-Raji cells lysates. Data are representative of three independent repeats.

Since HU-treatment increases *CXCL13* expression in Raji and HCT116 cells (Figure 2A, Supplementary Figure 2A, Figure 3E), we next examined whether MBD1 also regulates *CXCL13* expression under HU treatment. Compared to HU-treated shSCR-Raji cells, HU-treated shMBD1-Raji cells showed increased *CXCL13* expression (Figure 3E). The mean value of *CXCL13* mRNA in HU-treated shMBD1-Raji cells was 12.3 than the mean value of 10.6 in non-HU treated shMBD1 cells (Figure 3E-F). In addition, HU-treated shCTCF-Raji cells exhibited slight reduction in *CXCL13* levels than the HU-treated shSCR-Raji cells (Figure 3E). Moreover, HU-treated shCTCF+shMBD1-Raji cells exhibited almost equal levels of *CXCL13* expression as seen in HU-treated shMBD1-Raji cells, but these levels were greater than HU-treated shCTCF-Raji cells (Figure 3E). These results suggest that MBD1 plays a more important role in suppressing the *CXCL13* expression under HU-treatment (Figure 3E-F). These results also suggest that CTCF deficiency does not affect the upregulated levels of *CXCL13* in *MBD1*-deficient cells, suggesting a downstream role of CTCF in MBD1-dependent CXCL13-regulation (Figure 3E).

Indeed, *MBD1* overexpression in Raji cells led to a stronger reduction in *CXCL13* levels in response to HU treatment compared to non-HU-treated Raji cells (Figure 3F-G). Again, these analyses reveal a prominent role of MBD1 in suppression of *CXCL13* expression under the HU stress (Figure 3E-H). In contrast, overexpression of *BCL6*, a ubiquitously expressed protein in GC B cells [47], slightly reduced CXCL13 levels in either absence or presence of HU (Figure 3G, Supplementary Figure 3G). The expression of *BCL6* and *MBD1* plasmids was confirmed by western blotting in HEK293T cells (Supplementary Figure 3D, E).

Given that MBD1 binds to CpG-methylated DNA and promotes gene silencing [48], we next aimed to clarify the role of DNA methylation in *CXCL13* expression in other B-lymphomas and colon cancer cells (Figure 3H-I). From published datasets, we explored the effects of 5-Azacytidine (5-Aza), on *CXCL13* expression [49, 50]. 5-Aza is a chemotherapeutic drug that inhibits DNA CpG methylation [51]. Our analysis revealed that 5-Aza treatment in HT29 cells, a colon cancer cell line, induced *CXCL13* expression in the 5 μM and 10 μM 5-Aza-treated groups compared to the DMSO-treated groups [49] (Figure 3H, I). We also noted that 5-Aza treatment increased *CXCL13* and *CXCR5* expression in OCI-LY3 cells, a DLBCL cell type [50] (Figure 3I; GSE190319). These findings suggest the DNA-methylation dependent suppression of *CXCL13* expression. Finally, to test if cMYC, which is highly amplified in Burkitt’s lymphoma and the Raji cells due to translocation of the MYC locus to the Immunoglobulin locus, affects CXCL13 levels in Raji cells, we compared the CXCL13 levels in shSCR-Raji cells to shMYC#1-Raji and shMYC#2-Raji cells. The shMYC#1-Raji and shMYC#2-Raji cells showed a decrease in CXCL13 expression than shSCR-Raji cells (Figure 3J), indicating a positive role of c-MYC in promoting CXCL13 expression in Burkitt’s lymphoma cells. These results suggest that DNA methylation and MBD1 play a regulatory role in *CXCL13* expression, while cMYC positively regulates CXCL13 expression.

### CTCF and RNAPII enrichment on the *CXCL13* promoter upon HU-treatment in Raji cells

Since we observed a slight positive role of CTCF in *CXCL13* expression under non-HU treated condition (Figure 3E), we next attempted to understand the relationship between the pattern of CTCF binding on *CXCL13* promoter and its association with *CXCL13* expression. We explored the CTCF ChIP-seq data from 114 cell lines available on ENCODE (Figure 4A, Supplementary Figure 4A). We classified the CTCF-binding sites observed on the *CXCL13* locus in these 114 cell lines into three categories: site-1, site-2, and site-3 (Figure 4B). CTCF binding was observed at site-1 in 47 cell lines including the normal B and prostate cells (Figure 4A). The site-1 is located about 23 kb upstream of *CXCL13* TSS (Figure 4A, Supplementary Figure 4A). CTCF binding at site-2, consisting of two closer consecutive CTCF peaks, was observed in 101 out of 114 cell lines, including the normal B and prostate cells, known to be negative for CXCL13 expression (Figure 4A, Supplementary Figure 4A). The consistent pattern of CTCF binding at site-2 in 101 cell lines suggests a conserved binding of CTCF on the *CXCL13* locus (Figure 4A, Supplementary Figure 4A). Given that most cell types do not express the *CXCL13* naturally, we suggest that the conserved CTCF-binding on site-2 might be associated with suppression of *CXCL13* expression (Figure 4A). In contrast, only 8 cell types exhibited CTCF binding at site-3 (Figure 4A, supplementary Figure 4A), which spans between exons 2 and exon 3 of the *CXCL13* locus (Figure 4A, B). Interestingly, among these 8 cell types, 6 were of primary origin, while 2 cell lines, namely OCI-LY3 (DLBCL) and PC3 cell line (Prostate cancer), have been reported to exhibit *CXCL13* expression (Figure 4A, Supplementary Figure 4A) [52]. These findings suggest that CTCF binding at site-3 may be associated with an alternative conformation of the *CXCL13* promoter that could facilitate *CXCL13* transcription.

**Figure 4.**
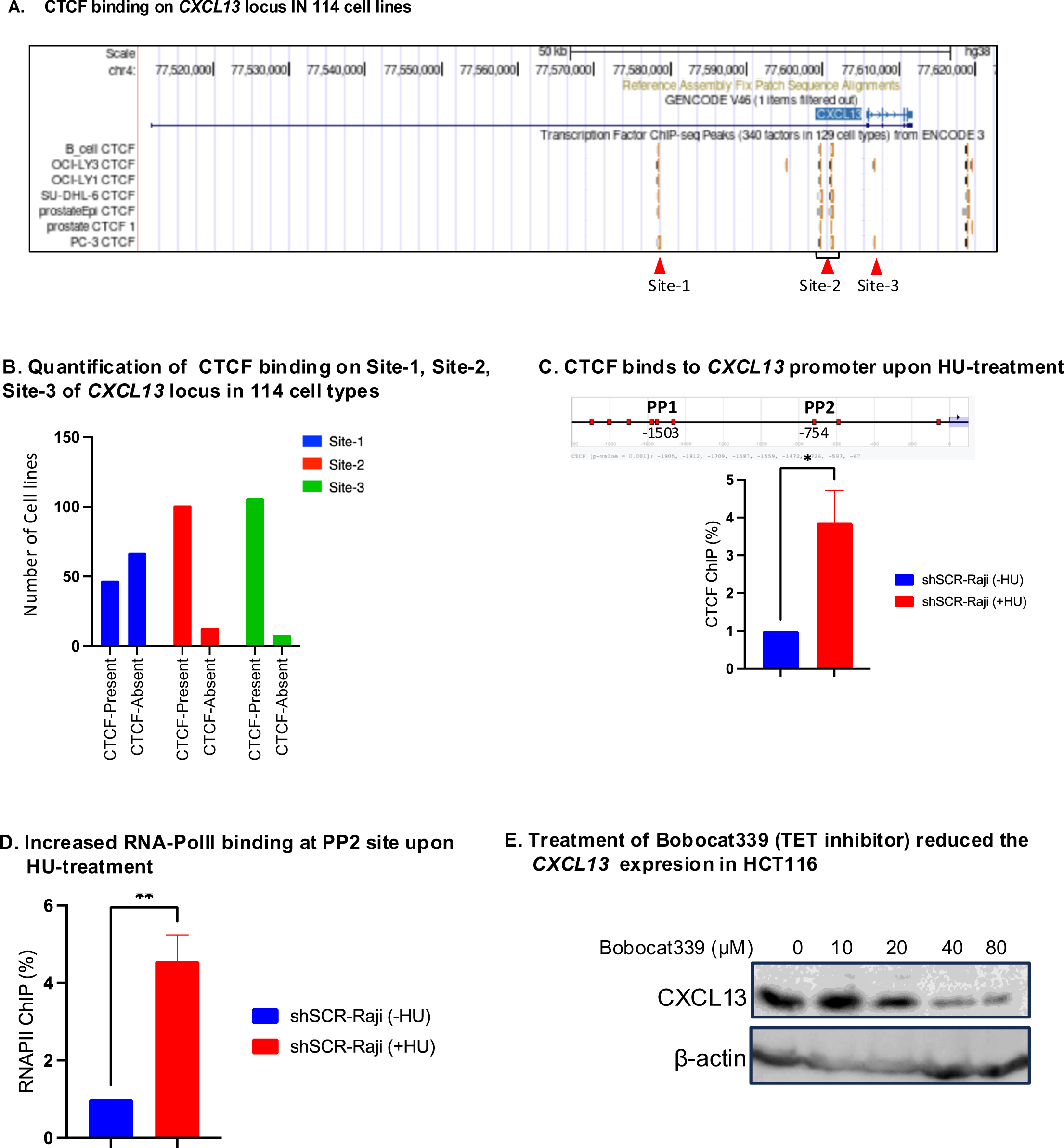
Enrichment of CTCF and RNAPII on *CXCL13* promoter in HU-treated Raji cells. (A) CTCF-ChIP-seq and distribution of CTCF binding on site-1, site-2 and site-3 on CXCL13 locus in normal human B cells, prostate cells and the OCI-LY3 and PC3 cell lines. The additional peak of CTCF on site-3 is located at position, chr4:77606576-77606790. Peak 1 sequence Position: chr4:77578302-77578777. Position of peak 2 from: chr4:77599430-77599754 OCI-LY3 and PC3 cell lines exhibit the *CXCL13* expression (B) Distribution of CTCF’s presence/absence on site-1, site-2 and site-3 in 114 cell types. Data represent the percentage of either presence or absence of CTCF peaks on site-1, site2 and sire-3 among 114 cell lines tested for CTCF ChIP-seq peaks. Loss of CTCF binding on either 1 or both peaks on Site-2 is counted as absence of CTCF binding. Data are derived from transcription factor ChIP-seq peaks from 129 cell lines on ENCODE from the UCSD genome browser (C) Schematic of CTCF putative binding sites on *CXCL13* locus at PP1 and PP2 sites located at −1503 and −597 upstream of the TSS respectively (D) CTCF ChIP-signals (%) on PP2 site in -/+HU treated Raji cells. Data are mean ± SEM for n=3. p= 0.0284 (unpaired t test) (E) RNAPII ChIP-signals (%) on PP2 site in -/+HU treated Raji cells. Data are mean ± SEM for n=3 p= 0.0058 (unpaired t test) (unpaired t test) (F) HCT116 cells were treated with DMSO and 10µM, 20µM, 40µM and 80µM concentrations of Bobocat-339 for 24 hours. After the treatment period, the cells were lysed, and proteins extracts were tested to assess the expression levels of CXCL13 by western blotting analysis.

To determine if CTCF redistribution occurs at the *CXCL13* promoter, we performed a CTCF-ChIP assay in HU-treated Raji cells (Figure 4C-D). HU treatment induces *CXCL13* expression (Figure 2A, 3E, Supplementary Figure 2A); therefore, altered CTCF binding at the *CXCL13* promoter may be observed under HU-treated conditions. *In silico* analysis predicted dense binding sites for CTCF between −1500 to −1800 and −597 and −726 on *CXCL13*-TSS (Figure 4I-J); these were designated as primary pair 1 (PP1) at −1503 bp (Figure 4I-J) and primer pair 2 (PP2) at −754 (Figure 4C-D). CTCF binding was not detected at the PP1 site, but CTCF binding was induced at the PP2 site in HU-treated Raji cells (Figure 4C-D). These suggests that HU-treatment leads to CTCF-redistribution on *CXCL13* locus in Raji cells (Figure 4C-D).

Since it is unclear whether CTCF redistribution is associated with *CXCL13* expression, we hypothesized that CTCF enrichment at the PP2 site might be linked to increased RNA Polymerase II (RNAPII) occupancy at the *CXCL13* locus (Figure 4E). To test this, we performed the RNAPII ChIP assay in HU-treated Raji cells (Figure 4E). Interestingly, we observed increased RNAPII occupancy at the PP2 site (Figure 4E). Based on these findings, we propose that CTCF recruitment to the *CXCL13* promoter leads to conformational changes of *CXCL13*-promoter, allowing the RNAPII and transcriptional machinery to access the *CXCL13* promoter and activate *CXCL13* expression under genotoxic stress.

Finally, considering the association between CTCF binding and TET2 dependent DNA-demethylation [53], we investigated whether TET1/2 inhibition affect the CXCL13 expression. Treatment with Bobcat-339, a potent TET1/2 inhibitor, reduced CXCL13 expression in HCT116 cells (Figure 4F). This suggests that TET1/2 enzymes promote CXCL13 expression in HCT116 cells. Although, we did not see an inhibitory effect of Bobocat-339 in Raji cells (Data not shown), this suggests a cell type specific effect of Bobocat-339 on CXCL13 expression. This may be possible that cell type specific genetic mutation may determine the TET1/2 activity on the *CXCL13* promoter. It is noteworthy that HCT116 cells harbors wild type p53 and Raji harbors two mutant p53. The p53 may interfere with TET1/2 activity on *CXCL13* promoter since Nultin-3a treatment induced the *CXCL13* transcription in Raji cells (Figure 2D). Overall, these results suggest a coordinated regulation among CTCF, TET enzymes, DNA methylation in determining the *CXCL13* expression in cancer and normal cells.

### Distal super-enhancer of *CCNG2* locus interacts with the *CXCL13* promoter in B-lymphoma

The human *CXCL13* is located on chromosome 4, and harbors 5 exons (Ensemble ID# ENST00000286758; CXCL13-201; Figure 5A, Supplementary 4B). Peripheral Blood Mononuclear Cells (PBMCs), including human B cells do not express *CXCL13* (Supplementary Figure 5A), suggesting a closed conformation of *CXCL13* promoter in most cell types. To elucidate the promoter conformation features of *CXCL13* expressing Raji cells, we assessed the binding of the RNAPII subunit POL2RA along the *CXCL13* locus using ENCODE ChIP-Seq data for Raji, as well as HepG2 and Ishikawa cells (Figure 5A). Strikingly, POL2RA ChIP-seq peaks were not observed in near vicinity of *CXCL13* promoter in Raji, HepG2 or Ishikawa cells (Figure 5A). The closest POL2RA peaks were found at −317,921 bp and −359,311 bp upstream of the *CXCL13* locus (Figure 5A), while the nearest downstream peak observed in Raji, HepG2, and Ishikawa cells was at +307,482 bp from the TSS (Figure 5A). Since we did not observe POL2RA peaks close to the *CXCL13*-TSS, we hypothesized that a distal enhancer may drive *CXCL13* expression in Cancer cells. To investigate this hypothesis, we explored the closest upstream and downstream regions co-occupied with POL2RA, histone H3K4me3, and the super-enhancer specific histone marks, the Histone H3K27-actyaltion (H3K27-Ac) and Histone H3K4me1 (Figure 5A). We analyzed the ChIP-seq data available at the ENCODE (UCSD database; Figure 5A-B), and identified three super-enhancers namely SE1, SE2 and SE3 (Figure 5A). SE1 is located between exon 2 and 3 of the *CCNG2* locus, while SE2 is downstream of *CCNG2*, and SE3 was present near the *CNOT6L* TSS (Figure 5A, C). The strongest POL2RA peaks were present at SE1 in Raji, HepG2 and Ishikawa cells, and the POL2RA peak on SE2 was only found in Raji cells but not in HepG2 or Ishikawa cells (Figure 5A). POL2RA occupancy was lower at SE2 than SE1 in Raji (Figure 5A). Given that SE1 harbors the highest enrichment of POL2RA signals (Figure 5A), we hypothesized that SE1 is a potential super-enhancer driving aberrant *CXCL13* expression in cancer cells. In mantle cell lymphoma, as demonstrated by HiC interaction data [54], there was an observed interaction between SE1 and the region flanking outside but nearer to *CXCL13* locus (Figure 5B). This interaction point further exhibited another interaction with the *CXCL13* locus (Figure 5B). Also, virtual circular chromosomal conformation capture (4C) from human spleen cells showed the interaction between region closer to SE1 and the *CXCL13* locus (Figure 5C) [55]. These suggest a possible interaction between the *CXCL13* promoter and the region near the *SE1* (Figure 5B,C). To test if SE1 interacts with *CXCL13* promoter, we performed chromosome conformation capture assay (3C) assay using Raji cells. We checked the interaction of SE1 with predicted *CXCL13*-TSS, located upstream of exon 1 [31] (Figure 5D-F). The 3C assay revealed an interaction between the *CXCL13* promoter and SE1 (Figure 5D, E). To confirm the interaction between SE-1 and the CXCL13 promoter, we performed the nested PCR from the purified PCR products using the primer FP2 and RP2 (Figure 5E, F). The nested PCR allows amplification from the first PCR containing 2069 bp product and generates a smaller band of 969 bps (Figure 5E, F). The presence of smaller band of 969 bps further confirmed the SE1-*CXCL13-TSS* interaction (Figure 5D-E). These results indicate that *CCNG2* super-enhancer interacts with *CXCL13*-TSS and induces the *CXCL13* expression in cancer cells. Since the SE1 is enriched with RNAPII, this interaction allows the access of RNAPII from SE1 to drive *CXCL13* transcription (Figure 5A,D-E). Moreover, 95 cell lines from ENCODE exhibited DNase-hypersensitivity sites and a higher value of GeneHancer score at SE1 (Figure 5A), suggesting the nature of open chromatin on SE1. Multispecies alignment of SE1 and SE3 revealed its conservation among the other vertebrates, suggesting a role for SE1 in developmentally programmed gene expression among vertebrates and its leaky access in cancer cells driving he *CXCL13* expression (Figure 5F).

**Figure 5.**
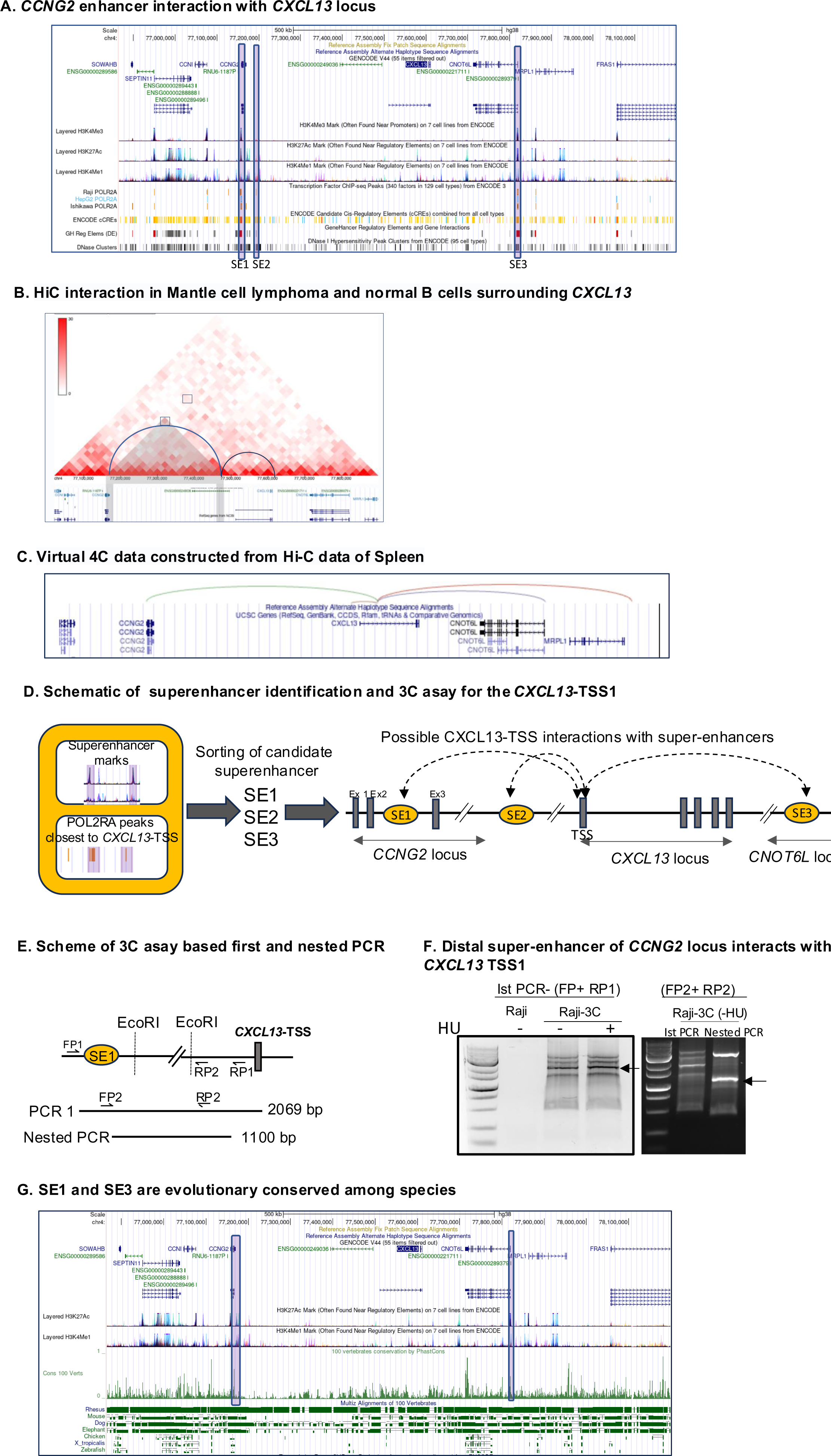
Distal super enhancer at *CCNG2* locus interacts with *CXCL13* promoter (A) (A) Genetic features of the *CXCL13* locus are depicted using the UCSD Genome Browser with Three super-enhancers surrounding the *CXCL13* locus, namely SE1, SE2, and SE3 are shown in maroon. Histone marks H3K4me3, H3K27Ac, and H3K4me1 are displayed from seven cell lines from the UCSD database. Super-enhancers highlighted in maroon and their overlapping with H3K27Ac and H3K4me1 signals. POL2RA ChIP-seq signals from ENCODE3 near the *CXCL13* locus are shown for Raji, HepG2, and Ishikawa cells. Comparison maps of micro-C and Hi-C interactions, GeneHancer regulatory elements, and DNase clusters of the *CXCL13* locus are presented from the UCSD database (B) HiC interaction data: Mantle cell lymphoma (left) and normal GC B cells (right). In the mantle cell lymphoma HiC map, the highlighted rectangle indicates an interaction between the *CCGN2* region and an area outside the *CXCL13* promoter, which also interacts with *CXCL13* promoter (C) Virtual Circular chromosomal conformation capture (4C) from spleen exhibiting the genomic interaction of *CXCL13* locus. Assembly hg19, resolution 40,000 base pairs (D) Schematic representation of the super-enhancer identification and their possible interaction with the *CXCL13* transcription start site (TSS). SE1, SE2, and SE3 exhibit overlapping signals of POL2RA, H3K27Ac, and H3K4me1. Schematic of SE1, SE2 and SE3 interaction with the super-enhancer located at *CCNG2* or *CNOT6L* (E) Schematic representation of 3C assay with EcoR1 digestion. Primer locations are shown for the detection of the first PCR and nested PCR products using primer pairs (FP1+RP1) and (FP2+RP2), respectively. The band lengths of the first PCR and nested PCR are 2069 and 1100 bp, respectively (F) Ethidium bromide staining of 3C assay PCR products. PCR performed with 1000 ng of purified genomic DNA from control Raji or 3C-processed (-/+HU) Raji cells. The nested PCR product for positive interaction between SE1 and *CXCL13*-TSS is shown with arrow in right (G) UCSD genome browser-based conservation of SE1, SE3 among 100 vertebrates by PhastCons and Multiz alignment is shown.

### *CXCL13* expression in B-lymphomas perturbs the CXCR5-migration

FDCs form CXCL13 gradient in the light zone (LZ) of GCs, which attracts CXCR5-positive GC B cells, and promotes their affinity maturation and selection [1, 9, 56]. Whether aberrant CXCL13 expression in GCDBLs serves as a programmed adaptation, promoting CXCR5-internalization and deprived LZ-migration is not known. Since peripheral blood mononuclear cell including B cells do no express *CXCL13* (Supplementary 5A), we hypothesized that *CXCL13* expression in GCDBLs may lead to CXCR5 receptor-internalization, reducing their migration towards external CXCL13. To test this, we compared CXCL13-induced migration of normal peripheral B cells (NPBCs) obtained from healthy donors with GCDBLs, Raji, FL-18, FL-218, and FL-318 (Figure 6A-C). NPBCs efficiently migrated towards recombinant human CXCL13 (rhCXCL13), while the migration index of multiple GCDBLs was significantly lower (Figure 6A). Strikingly, Raji cells exhibited robust migration towards recombinant human CXCL12 (rhCXCL12), a dark zone (DZ) chemokine (Figure 6B, C). This preference of Raji cells for rhCXCL12 over rhCXCL13 suggests GCDBLs confinement to the DZ and hinderance to LZ entry (Figure 6B, C). To investigate if internal CXCL13 disrupts CXCL13-mediated migration of GCDBLs towards an external CXCL13 gradient, we examined CXCL13-mediated migration of Raji cells transfected with lentivirus expressing CRISPR/Cas9 guide RNA (gRNA) targeting *CXCL13* exons 2 and 3 (referred to as *CXCL13*-KO Raji cells) (Figure 6D). Interestingly, bulk containing *CXCL13*-KO Raji cells exhibited a significant rescue in CXCL13-migration compared to Raji cells (Figure 6D). This suggests that internal CXCL13 disrupts migration of Raji cells and GCDBLs.

**Figure 6.**
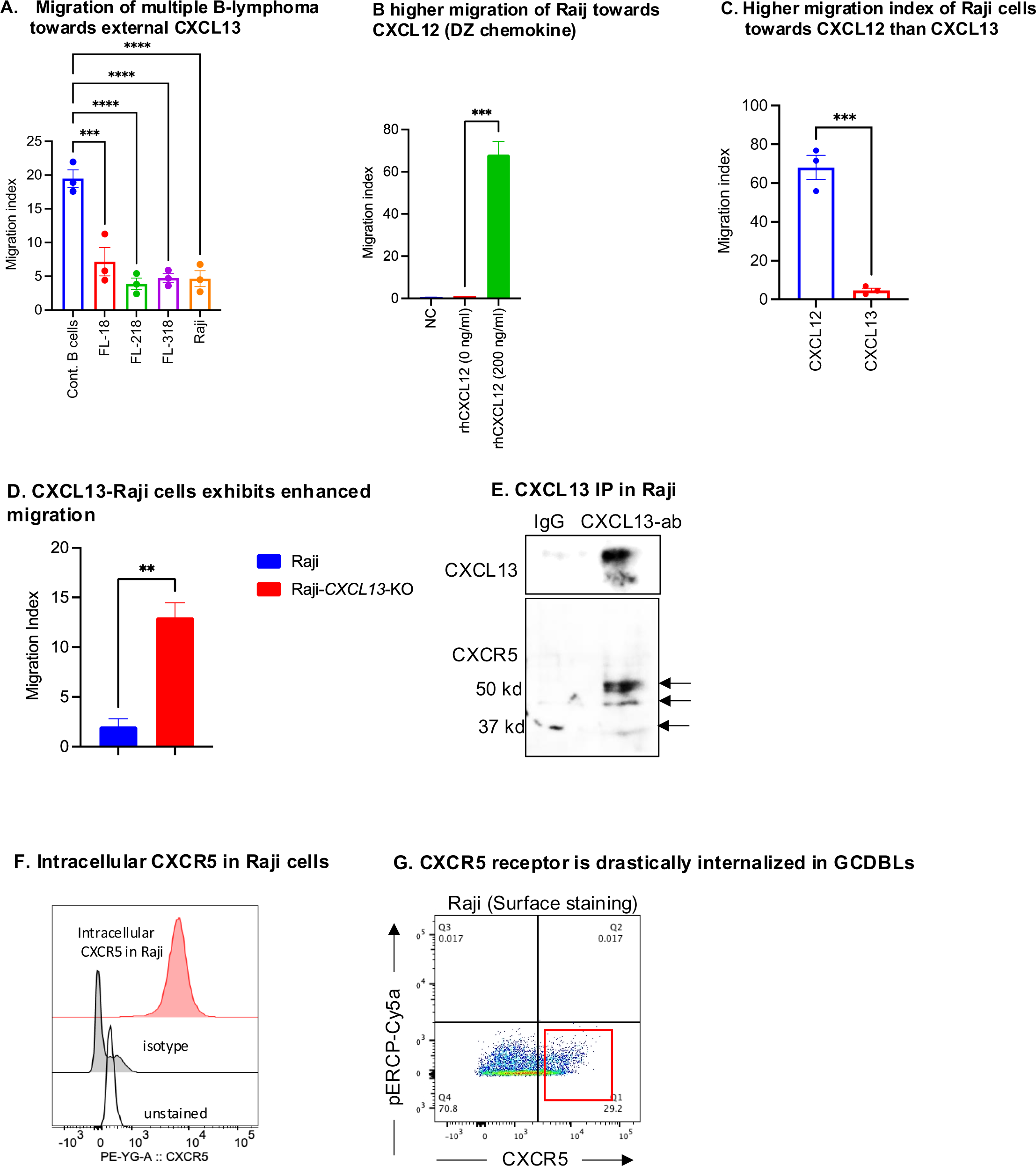
CXCL13 expression is human GCDBLs perturbs migration due to CXCR5-internalization. (A) Migration index of normal Human B and multiple Human B lymphoma lines Raji, FL-18, FL-518 and Granta-519 cell migration towards rhCXCL13 gradient. Data are mean ± SEM. P values for migration index were calculated using ordinary one-way Anova with multiple comparison. *p*= 0.002 for normal human B vs. FL-18, *p*<0.0001 for normal Human B vs FL-218, normal human B vs FL-318 and normal Human B vs Raji cells (B) Raji cells migrate efficiently towards rhCXCL12 (200 ng/ml). Data are mean ± SEM for n=3. NC= Negative control exhibiting the cell migrated to empty medium without the rhCXCL12 gradient (C) Raji cells exhibit enhanced migration towards rhCXCL12 compared to rhCXCL13 (n=3). The migration index of Raji cells towards rhCXCL12 was significantly higher than towards rhCXCL13 (p = 0.0006, unpaired Student’s t test) (D) Comparison of migration index between Raji cells and *CXCL13*-KO Raji cells towards rhCXCL13. *CXCL13*-KO Raji cells exhibit increased migration towards rhCXCL13 compared to control Raji cells (Data presented as mean ± SEM for n=3, p values calculated using unpaired t test) (E) Co-immunoprecipitation of CXCL13 and CXCR5 in Raji cells. Immunoprecipitation was performed using 100 μg of protein lysates from Raji cells with CXCL13 antibody (ab) or IgG antibody, followed by elution and Western blotting for CXCL13 and CXCR5 (F) Intracellular CXCR5 in Raji cells was detected using a CXCR5 antibody, followed by probing with a PE-conjugated secondary antibody. Unstained and isotype controls were used for flow cytometry staining. (G) Surface CXCR5 staining in Raji cells was performed with an APC-labelled CXCR5 antibody.

A plausible explanation of reduced migration in GCDBLs is CXCR5-internalization from interaction with internal CXCL13 ligand reduces its affinity towards external CXCL13. To test this, we performed co-immunoprecipitation (co-IP) of CXCL13 in Raji cells (Figure 6D). Elutes of CXCL13 co-IP exhibited distinct CXCR5 signals (Figure 6D), indicating a physical interaction between CXCL13 and CXCR5 in Raji cells (Figure 6D). This suggests that CXCR5-internalization takes place in Raji, thus B-lymphoma cells (Figure 6D). CXCR5-internalization was also confirmed by flow cytometry (FC) analysis: as intracellularly stained CXCR5 receptors were seen in 100% of Raji cells while only 29% of cells were positive for surface-CXCR5 (Figure 6F-G). These findings indicate internalization of surface-CXCR5 in Raji and possibly other GCDBL. Off note, to confirm that *CXCR5* coding sequence mutations were not the cause of reduced CXCL13 migration in Raji cells, we aligned Raji *CXCR5* with human *CXCR5* (Uniprot ID# P32302), revealing three SNPs (Supplementary Fig 6B). Despite these SNPs (Proline 39: CCC, Glycine 43: GGT, Threonine 338: ACC), however the resulting amino acids remained unchanged, confirming that Raji’s CXCR5 is identical to wild type CXCR5 (Supplementary Figure 6B). Therefore, alterations in the CXCR5 protein are not the cause of the altered migration of Raji cells towards CXCL13. Instead, our data suggests that the internalized CXCL13 caused the impaired migration of B-lymphoma cells (Supplementary Figure 6B).

### Accelerated cell cycle progression and suppression of p53-dependent transcripts by the CXCR5-CXCL13 axis in hematological and solid cancers

The CXCR5-CXCL13 axis activates intracellular signaling and confers chemotherapeutic resistance in *PTEN*-deficient cancers [32]. Given that *PTEN* is regulated by p53, this suggests a potential interference of the CXCR5-CXCL13 axis with p53 target genes. Conversely, studies have shown that p53 suppresses CXCR5 [57], indicating a complex interplay between p53 and the CXCR5-CXCL13 axis. Thus, the precise impact of the CXCR5-CXCL13 axis on p53 regulation remains poorly understood. To investigate the effects of CXCR5 overexpression on cancer cell survival, proliferation, chemotherapeutic resistance, and p53 regulation, we stably introduced the *pCMV-CXCR5-IRES-EGFP* plasmid into Raji cells *(Raji-CXCR5^oe^ hereafter,* Supplementary figure 7A, B). Notably, *Raji-CXCR5^oe^* cells displayed a marked increase in *CXCL13* levels (Supplementary Figure 7C), suggesting an activation of a positive feedback loop where CXCR5 signaling enhances *CXCL13* expression through autocrine loop (Supplementary Figure 7A-C). Despite this, *Raji-CXCR5^oe^* cells did not exhibit enhanced CXCL13-migration compared to Raji cells, indicating that increased CXCR5 levels did not drive CXCL13-mediated migration of GCDBLs, possibly due to increased intracellular CXCL13 levels and receptor internalization (Supplementary Figure 7D).

To assess the impact of the CXCR5-CXCL13 axis on cellular proliferation, we synchronized Raji and *Raji-CXCR5^oe^* cells using HU treatment. After releasing the synchronized cells (Figure 7A), we monitored the live cell number (Figure 7A, B). Interestingly, *Raji-CXCR5^oe^* cells demonstrated significantly higher proliferation rates than Raji cells (Figure 7B). We also observed elevated *JUN* levels in *Raji-CXCR5^oe^* cells, a downstream target of CXCR5-CXCL13 signaling (Figure 7C), particularly in the presence of bleomycin or a combination of bleomycin and Nutlin-3a (Fig 7C). Compared to control Raji cells, *Raji-CXCR5^oe^* cells showed increased levels of *CCR7* (Figure 7C), a GPCR known to promote cancers metastasis [58]. Treatment with bleomycin, Nutlin-3a, or the combination of both led to increased *CCR7* levels in both Raji and *Raji-CXCR5^oe^* cells (Figure 7C). However, the levels of *CCR7* were notably higher in *Raji-CXCR5^oe^* cells within their respective treatment groups compared to Raji cells (Figure 7C). In contrast *Raji-CXCR5^oe^* cells exhibited an overall reduction in *TP53* transcript levels compared to Raji cells (Figure 7D). *TP53* transcripts were also low in individual or simultaneously treated groups with bleomycin and Nutlin-3a (Figure 7D) in the *Raji-CXCR5^oe^* than Raji cells, indicating a potential interference of the CXCR5-CXCL13 axis in the p53 levels (Figure 7D).

**Figure 7.**
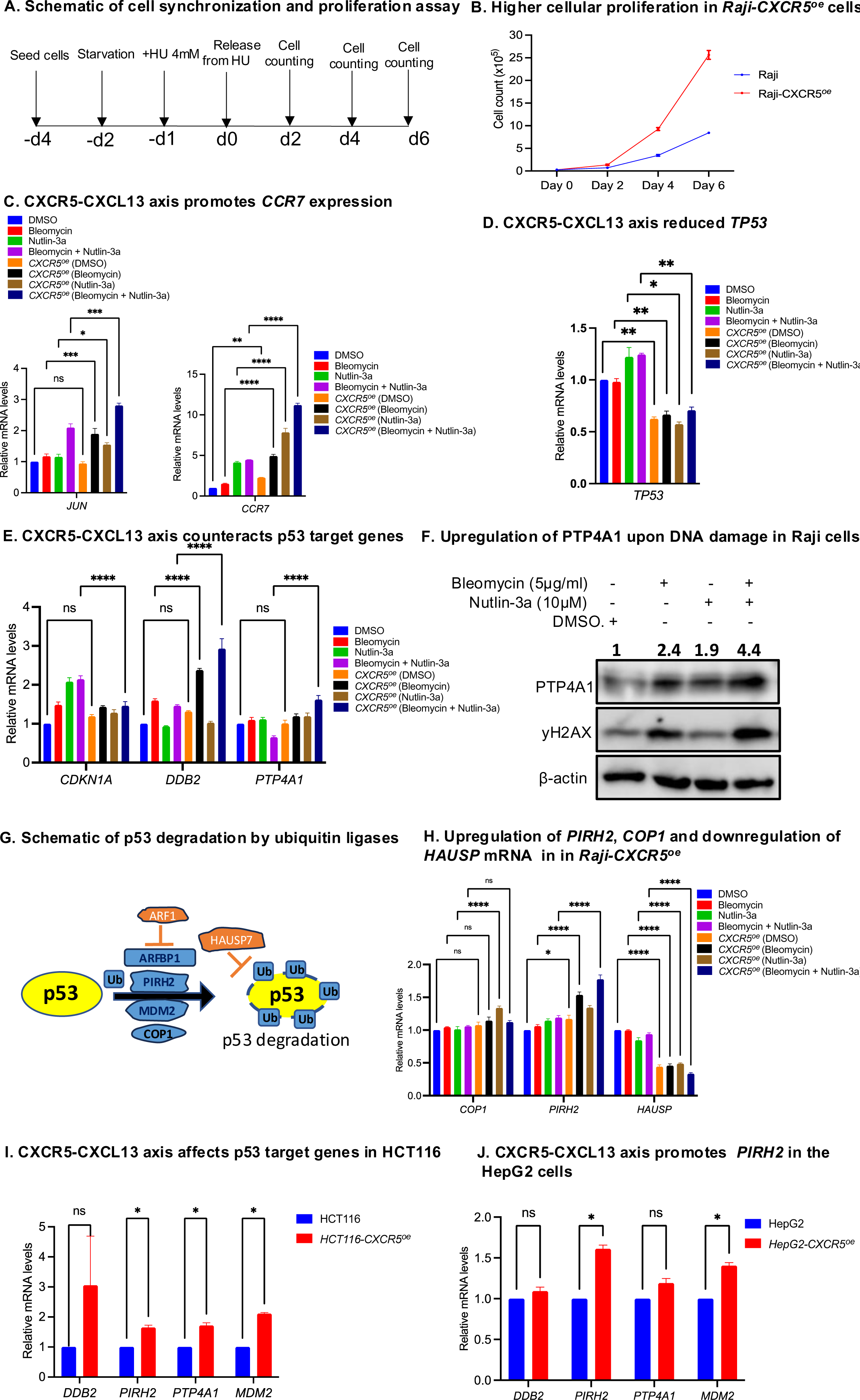
CXCR5 Modulates Cell Cycle Progression and Overrides p53 response. (A) Schematic representation of cell synchronization by HU-treatment of Raji and *Raji-CXCR5°*^e^ cells (B) Enhanced proliferation in *Raji-CXCR5^oe^* cells compared to Raji cells. Cell proliferation assessed on days 0, 2, 4, and 6 with data from four replicates, plotted as multiples of (1*10^5^) cells on the X-axis (C) Treatment of Raji and *Raji-CXCR5^oe^*cells with Bleomycin (5μg/ml), Nutlin-3a (10μM), or their combination for 24 hours and qRT-PCR quantification of *JUN* and *CCR7* transcript levels in *Raji-CXCR5^oe^* cells compared to Raji cells. *JUN* transcript is shown as downstream positive marker gene induced from CXCR5-CXCL13 signaling. p= 0.0002 for Raji (Bleomycin) vs *Raji-CXCR5^oe^* (Bleomycin): p= 0.0336 for Raji (Nutlin-3a) vs *Raji-CXCR5^oe^* (Nutlin-3a): p= 0.0002 in Raji (Nutlin-3a+bleomycin) vs *Raji-CXCR5^oe^*(Nutlin 3a+bleomycin) groups. Šídák’s multiple comparisons test. For *CCR7* transcripts; p= 0.0022 for for Raji (DMSO) vs *Raji-CXCR5^oe^*(DMSO) and p<0.001 for Raji vs *Raji-CXCR5^oe^* in all three respective treatment groups. Šídák’s multiple comparisons test. Data presented as mean ± SEM for n=3. (D) qRT-PCR quantification of *TP53* transcript levels in *Raji-CXCR5^oe^* cells compared to Raji cells. p =0.0025 in Raji (DMSO) vs *Raji-CXCR5^oe^*(DMSO), p=0.0064 Raji (bleomycin) vs *Raji-CXCR5^oe^*(Bleomycin), p= 0.0114 for Raji (Nutlin-3a) vs *Raji-CXCR5^oe^*(Nutlin-3a) and p= 0.0052 for Raji (bleomycin+Nutlin-3a) vs *Raji-CXCR5^oe^*(Bleomycin+Nutlin-3a) cells. p values are derivatives of t test. Data presented as mean ± SEM for n=3 (E) Treatment of Raji/*Raji-CXCR5^oe^*cells with Bleomycin (5μg/ml), Nutlin-3a (10μM), or their combination and mRNA quantification of p53 target genes, *CDKN1A*, *DDB2* and *PTP4A1*. p <0.0001 for *CDKN1A*, *DDB2*, *PTP4A1* in Raji (Bleomycin+Nutlin-3a) vs *Raji-CXCR5^oe^*(Bleomycin +Nutlin-3a). p values are derived with Tukey’s multiple comparison test. Data presented as mean ± SEM for n=3 (F) Treatment of Raji cells with Bleomycin (5μg/ml), Nutlin-3a (10μM) for 24 hr and Western blotting analysis of PTP4A1 and γH2AX using whole cell lysates. PTP4A1 levels were increased after bleomycin and Nutlin-3a treatment of Raji cells. Values above blot indicate densitometric qualification of PTP4A1 normalized to β-actin levels. Data are representative of three independent replicates (G) Schematic representation of enzymes regulating the ubiquitination/de-ubiquitylation of p53 (H) Treatment of Raji/*Raji-CXCR5^oe^* cells with Bleomycin (5μg/ml), Nutlin-3a (10μM), or their combination and qRT-PCR quantification of *COP1*, *PIRH2* and *HAUSP* in *Raji-CXCR5^oe^*cells compared to Raji cells. p< <0.0001 for *COP1* in Raji (Nutlin-3a) vs *Raji-CXCR5^oe^* (Nutlin-3a). p= 0. 0.0118 for *PIRH2* in Raji (DMSO) vs *Raji-CXCR5^oe^*(DMSO), p<0.0001for *PIRH2* in Raji (Bleomycin) vs *Raji-CXCR5^oe^*(Bleomycin). p <0.0001 for *HAUSP* in all respective treatment groups of Raji vs *Raji-CXCR5^oe^* cells. Tukey’s multiple comparisons test. Data presented as mean ± SEM for n=3 (I) HCT116 and *HCT116-CXCR5^oe^* cells utilized for qRT-PCR analysis for *DDB2*, *MDM2*, *PIRH2*, and *PTP4A1* mRNA levels. p= 0.276680 (*DDB2*), p= 0.000845 (*PIRH2*), p= 0.001291 (PTP4A1), p= 0.000005 (MDM2) in HCT116 vs *HCT116-CXCR5^oe^* cells. Unpaired t test. Data presented as mean ± SEM for n=3. (J) HepG2 and *HepG2-CXCR5^oe^* cells utilized for qRT-PCR analysis of for *DDB2*, *MDM2*, *PIRH2*, and *PTP4A1* mRNA levels. p= 0.126974 (*DDB2*), p= 0.00017 (*PIRH2*), p= 0.026746 (PTP4A1), p= 0.000404 (MDM2) in HepG2 vs *HepG2-CXCR5^oe^* cells. Unpaired t test. Data presented as mean ± SEM for n=3.

The transcriptional targets of p53 play important roles in cell cycle regulation, therefore we measured mRNA levels of *p53* target genes *CDKN1A* (encodes the p21), *DDB2* (encodes a DNA-repair factor) and *PTP4A1*(encodes for a protein phosphatase required for mitotic entry) between Raji and *Raji-CXCR5^oe^* cells (Figure 7E). Following treatment with bleomycin, Nutlin-3a, or their combination to induce p53 activation, we observed an induction of *CDKN1A* and *DDB2* levels in Raji cells (Figure 7E). Interestingly, *Raji-CXCR5^oe^* cells exhibited a lower induction of *CDKN1A* in response to combined treatment with bleomycin and Nutlin-3a compared to Raji cells treated with bleomycin and Nutlin-3a (Figure 7E). Conversely, *DDB2* levels were notably higher in *Raji-CXCR5^oe^*cells, particularly in the groups either treated with bleomycin or co-treated with bleomycin and Nutlin-3a, compared to the corresponding treatment groups of Raji cells (Figure 7E). Furthermore, the expression of *PTP4A1* was notably increased in *Raji-CXCR5^oe^*cells upon combined treatment with bleomycin and Nutlin-3a, as compared to its decreased expression in Raji cells with similar treatment conditions (Figure 7E). This indicates that the CXCR5-CXCL13 axis can impede the p53-mediated regulation of *PTP4A1, DDB2* and *CDKN1A* (Figure 7D). Since Raji cells exhibited DNA-damage-induced expression of CXCL13 and CXCR5 (Figure 2C, D), we investigated whether this upregulation of CXCR5 and CXCL13 could enhance PTP4A1 levels without the need for CXCR5 overexpression. Our observations revealed that treatment of Raji cells with Bleomycin and Nutlin-3a led to nearly a twofold increase in PTP4A1 protein levels (Figure 7F). Moreover, this enhancement of PTP4A1 was further intensified, reaching up to a fourfold increase, upon combined treatment of Raji cells with Bleomycin and Nutlin-3a (Figure 7F). These findings confirm that the chemotherapeutic-induced CXCR5-CXCL13 signaling can stimulate PTP4A1 expression, potentially promoting cell cycle proliferation and contributing to chemotherapeutic resistance (Figure 7F). While co-treatment of Raji cells with bleomycin and Nutlin-3a led to a decrease in *PTP4A1* mRNA levels, there was no significant impact on protein levels. This implies that additional regulatory mechanisms may regulate PTP4A1 in Raji cells following the combined treatment with bleomycin and Nutlin-3a (Figure 7F).

Levels of p53 are regulated by ubiquitin ligases MDM2, COP1, ARF1BP1 (HUWE1) and PIRH2,(Figure 7G) [59, 60]; in turn p53 can be stabilized by HAUSP-mediated de-ubiquitination of p53 itself, ARF inhibition of ARFBP1 ubiquitin ligase activity, or ARF binding and sequestration of MDM2 [61]. To assess how CXCR5 regulates p53, we first measured expression of these ubiquitin ligases in *Raji-CXCR5^oe^* cells (Figure 7G, H). Compared to Raji cells, *Raji-CXCR5^oe^* cells exhibited elevated levels of *PIRH2* (*RCHY1*), while no significant increase was observed in *COP1* levels (Figure 7H). However, *COP1* levels were induced in Nutlin-3a-treated *Raji-CXCR5^oe^* cells compared to Raji cells treated with Nutlin-3a (Figure 7H). In contrast, *PIRH2* levels were significantly elevated in the DMSO-treated, bleomycin-treated or the combination of bleomycin and Nutlin-3a treated groups of *Raji-CXCR5^oe^* cells compared to respective groups in Raji cells (Figure 7H).The levels of ARF1 were decreased in *Raji-CXCR5^oe^*cells than Raji cells (Supplementary Figure 7E). Notably, *ARFBP1* and *MDM2* levels were also reduced in *Raji-CXCR5^oe^* cells than Raji cells (Supplementary Figure 7E). The levels of HAUSP, which stabilizes p53 by promoting its deubiquitylation, were significantly reduced in *Raji-CXCR5^oe^* cells (Figure 7H). In contrast, *Raji-CXCR5^oe^* cells treated with bleomycin, Nutlin-3a or a combination of both showed a significant reduction in the *HAUSP* levels compared to respective treatment grouped in Raji cells (Figure 7 H). These suggests a potential role for CXCR5-CXCL13 signaling in destabilizing p53 through decreased HAUSP and increased PIRH2 levels (Figure 7H). These findings suggest that the CXCR5-CXCL13 axis may modulate p53 levels by disrupting the balance of its ubiquitination and deubiquitinating enzymes (Figure 7H).

Raji cells harbor mutant p53 proteins, so we further analyzed the effect of CXCR5-CXCL13 axis on p53-target genes using the HCT116 and HepG2 cells, the COAD and LIHC cell lines respectively (Figure 7I, J). HCT116 and HepG2 harbor wildtype p53, which allows us to examine the effect of CXCR5-CXCL13 on p53 target genes accurately. HCT116 and HepG2 cells exhibit *CXCL13* expression, therefore we transiently overexpressed the *CXCR5* in HCT116 and HepG2 cells to activate the CXCR5-CXCL13 axis (*HCT116-CXCR5^oe^*and *HepG2-CXCR5^oe^* hereafter). Interestingly, *HCT116-CXCR5^oe^*cells exhibited significant upregulation of *PIRH2*, *MDM2*, and *PTP4A1* than HCT116 cells (Figure 7I). Furthermore, *HepG2-CXCR5^oe^* cells also exhibited significant upregulation of *PIRH2* and *MDM2* than HepG2 cells (Figure 7J). The most induced genes were *PIRH2*, *PTP4A1*, and MDM2 in Raji, HCT116 and HepG2 after CXCR5 overexpression (Figure 7E-J). Collectively, these results suggest that the CXCR5-CXCL13 axis promotes the expression of p53 specific ubiquitin ligases, *PIRH2* and *PTP4A1* in lymphoid and solid cancers suggesting a role of CXCR5-CXCL13 axis in destabilizing the p53 levels and promoting cell cycle progression (Figure 7B, H). Collectively, these results suggest that CXCR5-CXCL13 axis impairs the p53 axis, promoting the tumor proliferation.

## Discussion

### Regulation of *CXCL13* expression by DNA methylation, MBD1, CTCF and TET enzymes

The observed inhibitory effects of 5-Aza treatment and MBD1 on *CXCL13* expression in colorectal cancer, DLBCL, and Raji cells (Figure 3E-I) are consistent with previous findings highlighting the suppressive role of the DNA methylation axis on gene expression [62–64]. Furthermore, the influence of TET enzymes and CTCF on *CXCL13* expression (Figures 3E, 4A-E) suggests a complex interplay involving DNA methylation, MBD1, TET enzymes, and CTCF in regulating *CXCL13* transcription. However, the stepwise mechanisms by which DNA methylation, MBD1 regulation, TET activity, and CTCF regulation affect *CXCL13* expression require further investigation. We could clarify that under the DNA-replication stress, MBD1 works upstream of CTCF in *CXCL13* regulation since the double knockdown of *MBD1* and *CTCF* induced the similar level of *CXCL13* levels as seen in single knockdown of MBD1 (Figure 3E). On the other hand, CTCF seems to also affect *CXCL13* expression under normal conditions (Figure 3E), suggesting a partially non-overlapping role of CTCF and MBD1 on *CXCL13* transcription (Figure 3E). Future studies should investigate whether TET enzymes affect CTCF distribution and MBD1 regulation at the *CXCL13* locus in cancer cells, reported previously [53, 65, 66].

### ATR-signaling, TET activity and *CXCL13* expression

We observed that DNA damage and ATR signaling promote CXCL13 expression (Figure 2A, Supplementary Figure 2A, Figure 2H). This suggests that DNA damage and ATR signaling may alleviate the suppressive effects of DNA methylation and MBD1 on *CXCL13* transcription. This aligns with ATR’s role in TET3-dependent DNA demethylation and gene expression [67]. TET inhibition reduced CXCL13 expression in HCT116 cells (Supplementary Figure 2A), suggesting that DNA damage and ATR signaling may recruit TET enzymes to the *CXCL13* locus, leading to DNA hypomethylation and induction of *CXCL13* transcription. A rational for ATR-dependent *CXCL13* transcription could be in activating the autocrine signaling in CXCR5+ cancer cells, and accelerated DNA repair. This is supported by observation that Raji cells treated with 20mM HU, which have higher CXCL13 levels, showed lower γH2AX signals compared to those treated with 10mM HU (Figure 2A, B). This suggests a novel link between ATR signaling and TET-dependent CXCL13 expression in DNA repair.

### CTCF, enhancer hijacking and RNAPII enrichment on the *CXCL13* promoter

Using 3C assay, we identified that a distal super-enhancer (SE1) at the *CCNG2* locus interacts with the *CXCL13* transcription start site (TSS) in Raji cells (Figure 5B-D). Evidence from Raji, HepG2, and Ishikawa cells shows that the *CXCL13*-TSS lacks RNAPII signals at least 300 kb upstream and downstream (Figure 5A). This suggests that SE1 interaction may facilitate transcriptional machinery access to *CXCL13*-TSS in cancer cells (Figure 4E, Figure 5A, C). It is possible that the interaction between SE1 and the *CXCL13* promoter may be regulated by TET enzymes and CTCF. Supporting this, HU treatment, which induces *CXCL13* expression (Figure 2a, 3E) in Raji cells, leads to CTCF and RNAPII enrichment at the CXCL13 promoter (Figure 4D-E). This indicates coordinated localization of CTCF and RNAPII under HU stress on *CXCL13* promoter. Additionally, CTCF peaks at the *CXCL13* promoter site-3 are found in CXCL13 positive OCI-LY3 and PC3 cells but not in normal B cells or hepatocytes (Figure 4A-D), suggesting CTCF distribution may influence the inactive state of the *CXCL13* promoter. Further studies are needed to determine whether the interaction between SE1 and the CXCL13 promoter depends on CTCF redistribution, potentially altering chromosomal conformation within or beyond the associated topologically associated domain, thereby facilitating RNAPII access to the *CXCL13* locus. These findings align with the established interplay among TET enzymes, DNA hypomethylation, CTCF binding, and chromosomal conformation changes during development [53, 65, 66].

### Conserved CTCF binding on *CXCL13* locus and transcriptional suppression

The *CXCL13* locus exhibits a consistent and conserved pattern of CTCF binding on site-2 in the 101 out of 114 examined cell lines (Figure 4A,B). Based on ENCODE ChIP-seq data sets CTCF binding at site-2 of the *CXCL13* locus overlaps with RAD21 and SMC3 peaks in the normal hepatocytes. The CTCF-RAD21-SMC3 may block general transcription elongation, (Nanavaty, 2020, 32333838), suggesting a suppressive role of site-2 in *CXCL13* expression. It may be possible that cohesin defective cells may exhibit the *CXCL13* overexpression in few cancer types. ENCODE data suggest the loss of RAD21 in one sample of HepG2 in site 2, suggesting that RAD21 and cohesin loss may promote the CXCL13 expression in cancer cells.

### Physiological elimination of B-lymphomas during humoral immune response

CXCL13 expression in B-lymphomas may represent a programmed adaptation that restricts their migration to the LZ by CXCR5-internalization and confines them to the DZ. This restriction could potentially limit survival feedback signaling from FDCs, leading to the GCDBLs elimination. The orchestrated *CXCL13* expression via CTCF and altered chromosome conformation (Figure 3A-D, Figure 5A-D) might hinder the LZ-migration and fate of GC B cells. Moreover, Burkitt’s lymphomas, and myeloma exhibit translocations between the immunoglobulin heavy and *MYC* loci [68–70], resulting in MYC overexpression. MYC is required for LZ entry of GC B cells [71]. The amplified *MYC*, leading to CXCL13 overexpression, may simultaneously prevent the LZ entry of GCDBLs, suggesting a dual role for MYC in regulating the fate of germinal center B cells. This indicates a unique adaptation within GCDBLs, where MYC and oncogenic factors like BCL6 may paradoxically serve tumor-suppressive roles by preventing the exit of GCDBLs from GCs [72]. These regulations assist the robust immune response by facilitating the elimination of GCDBLs and promoting the generation of high-affinity antibodies. Furthermore, internalized CXCR5 may undergo proteasomal degradation in GCDBLs and needs to be tested in future studies.

### Chemotherapeutic resistance

The inducibility of CXCL13 and CXCR5 by DNA damage-inducing chemotherapeutics highlights potential concerns on resistance caused by CXCR5-CXCL13 axis in cancer cells (Figure 2A-D, Supplementary Figure 2A). We show that the CXCR5-CXCL13 axis can counteract p53 levels by increasing expression of the ubiquitin ligase *PIRH2* and reducing the expressions of its de-ubiquitylation enzyme, *HAUSP* (Figure 7H-I). CXCR5-CXCL13 signaling also counteracted the p53 target genes *PTP4A1*, *DDB2* and *CCR7* (Figure 7D-I), which could potentially promote cell cycle progression, DNA repair and metastasis. It will be important to see if *PTP4A1*/*CCR7/DDB2* knockdown in CXCR5 and CXCL13 co-expressing cancers perturbs tumor growth in xenograft models. *PTP4A1* and *CCR7* could be novel markers exhibiting chemotherapeutic-resistance in the cancer exhibiting the CXCR5-CXCL13 signaling.

## Declarations

### Ethical Approval

The use of human B cells/ cell lines was approved as per Institutional ethical board.

### Authors’ contributions

SKG designed the original hypothesis, performed experiments, and analyzed data. PM discussed the results and provided constructive feedback. SKG wrote the manuscript. JHB critically read the manuscript and provided constructive feedback. All authors read and agreed to manuscript.

### Availability of data and materials

All data needed to evaluate the conclusions in the paper are present in the main text and the supplementary materials.

## Acknowledgment

The authors acknowledge TCGA and the ENCODE Consortium for generating data used in this study. We thank Dr. Katsuya Sakai for anti-goat IgG-HRP antibody. We thank Dr Susumu Kohno for gifting shMYC plasmids. We thank Dr. Atsuyasu Sato for the discussion. We thank Dr. Atsushi Hirao for providing the THP1 cells and Dr. Kazuhiro Murakami for useful discussions and providing the HCT116 and LoVo cell lines, we thank Dr. Y. Endo for HepG2 cells. We thank Dr. Takeshi Suzuki for the supporting SKG’s research at Kanazawa University.

## Limitations of the study: Limitations of the study

In this study, we have identified and characterized a potential mechanism that inhibits *CXCL13* expression in adherent cells, as CXCL13 is typically expressed in certain immune cells. Our findings suggest that the suppression of *CXCL13* expression involves DNA methylation factors and a redistribution of CTCF on the *CXCL13* promoter, leading to a conformational change that transitions the promoter to a transcriptionally active state. However, further investigation is required to elucidate whether DNA methylation and TET activity coordinate these changes, and how the loss of DNA methylation and CTCF redistribution influences this altered chromosomal conformation and RNAPII enrichment on *CXCL13* promoter to activate transcription. Additionally, the CTCF boundaries that overlap with RAD21 and SMC3 seem to play a regulatory role in the polyadenylation of *CXCL13* mRNA, potentially resulting in various *CXCL13* transcript forms. Therefore, transcriptional elongation may be perturbed by the RAD21/CTCF/SMC3 sites on intron 2, producing alternatively spliced *CXCL13* transcripts. We have not addressed the alternative splicing of *CXCL13* in this manuscript. Our study does not conclusively distinguish the forms of the CXCL13 protein reported. This is also to note that the CXCL13 molecular weight appears higher than the predicted 13 kDa band. This will also help clarify whether intracellular CXCL13 expression contributes to CXCR5 receptor internalization. Attempts to generate viable CXCL13-knockout cell lines other than in Raji cells may be necessary to resolve this concern. A secretory form of CXCL13 may arise from the reported variant, which could impact on CXCR5-CXCL13 signaling in autocrine and paracrine mode, but its appearance may not be visible on WB due to secretion. These limitations warrant further investigation in future studies.

---

**Supplementary Figure 1.**
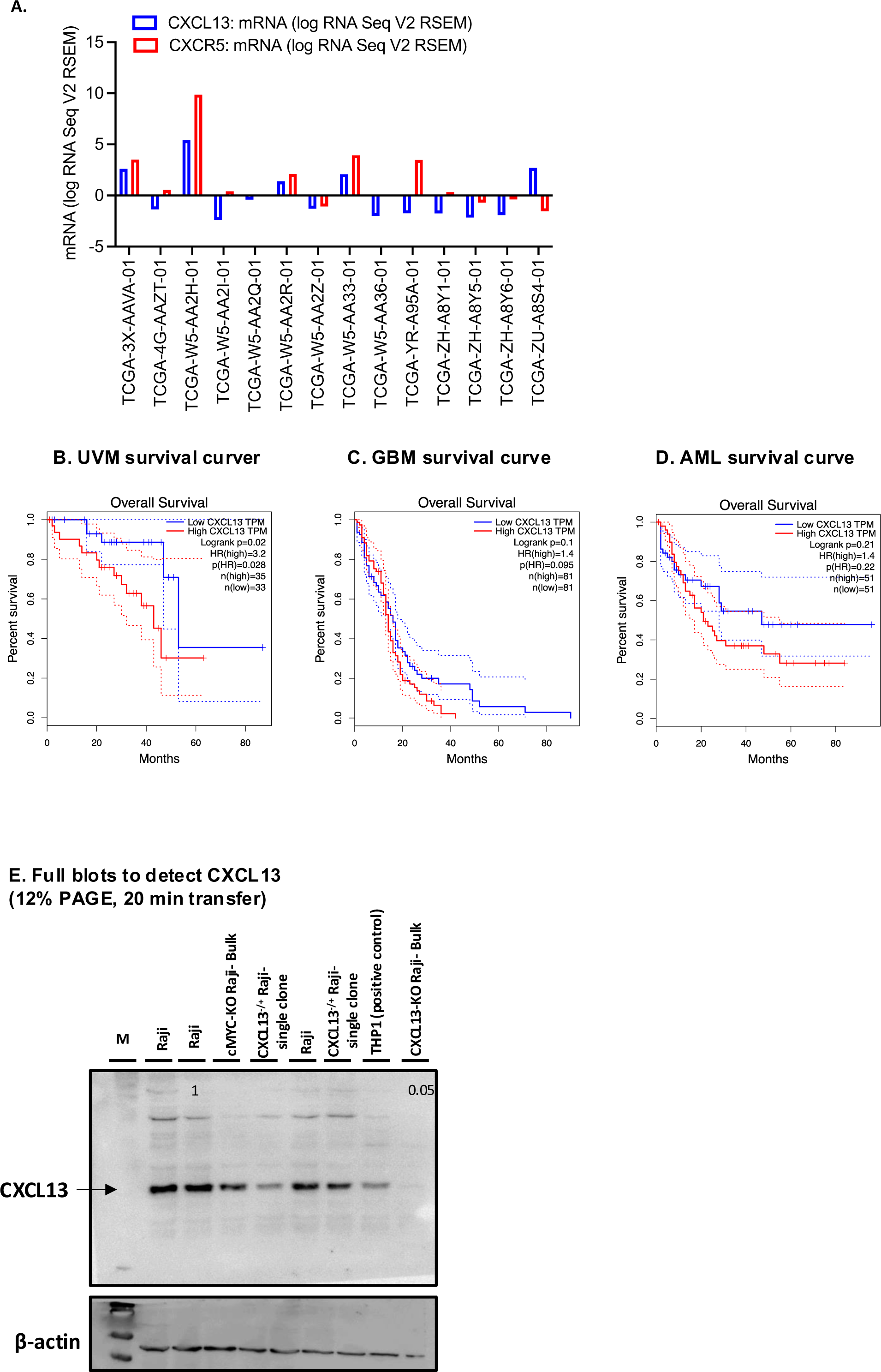
(A) Comparison of *CXCR5* and *CXCL13* expression in 7 CHOL samples revealed co-expression of both *CXCR5* and *CXCL13* in 4 samples. Co-expression indicates a possible autocrine signaling in these cancers. (B-D) Overall survival analysis from *GEPIA* (Gene Expression Profiling Interactive Analysis) indicates that groups with low *CXCL13* TPM and high *CXCL13* TPM for Uveal Melanoma (UVM), Glioblastoma (GBM), and Acute Myeloid Leukemia (AML) demonstrate varying survival outcomes. The *CXCL13*-high TPM groups exhibit reduced survival rates in these cancer types (E) Detection of CXCL13 in cell lysates of Raji cells, CRISPR/Cas9 transfected Raji cells with guide RNA targeting *MYC*/*CXCL13* cells. The THP1 cell line serves as a positive control, expressing CXCL13. Antibody:Novus#AF801. Cell lysates of CRISPR/Cas9 transfected Raji cells were prepared after transfection with CRISPR/Cas9 plasmid targeting exon 2 and 3 of *CXCL13* locus, followed by culture for 2-days prior to sorting of transfected cells. Sorted cells were subsequently cultured and single cell clones were grown until confluency and viability. The single-cell clones (*CXCL13^-/+^*) were then subjected to Western blot analysis. The SDS-PAGE gel concentration used was 12%, and the transfer was conducted for 20 minutes. Homozygous clones *(CXCL13^-/-^)* of Raji cells could not grow faster and maintained in passage. The Western blot was conducted using a bulk of sorted cells that were positively transfected with CRISPR/Cas9, following their culture until viability was maintained. CXCL13 appeared around 22 kilodaltons (kDa), which aligns with the weight of CXCL13 in THP1 cells—a positive control cell line expressing CXCL13 and is reduced in two independent *CXCL13^+/-^* clones of Raji cells.

**Supplementary Figure 2.**
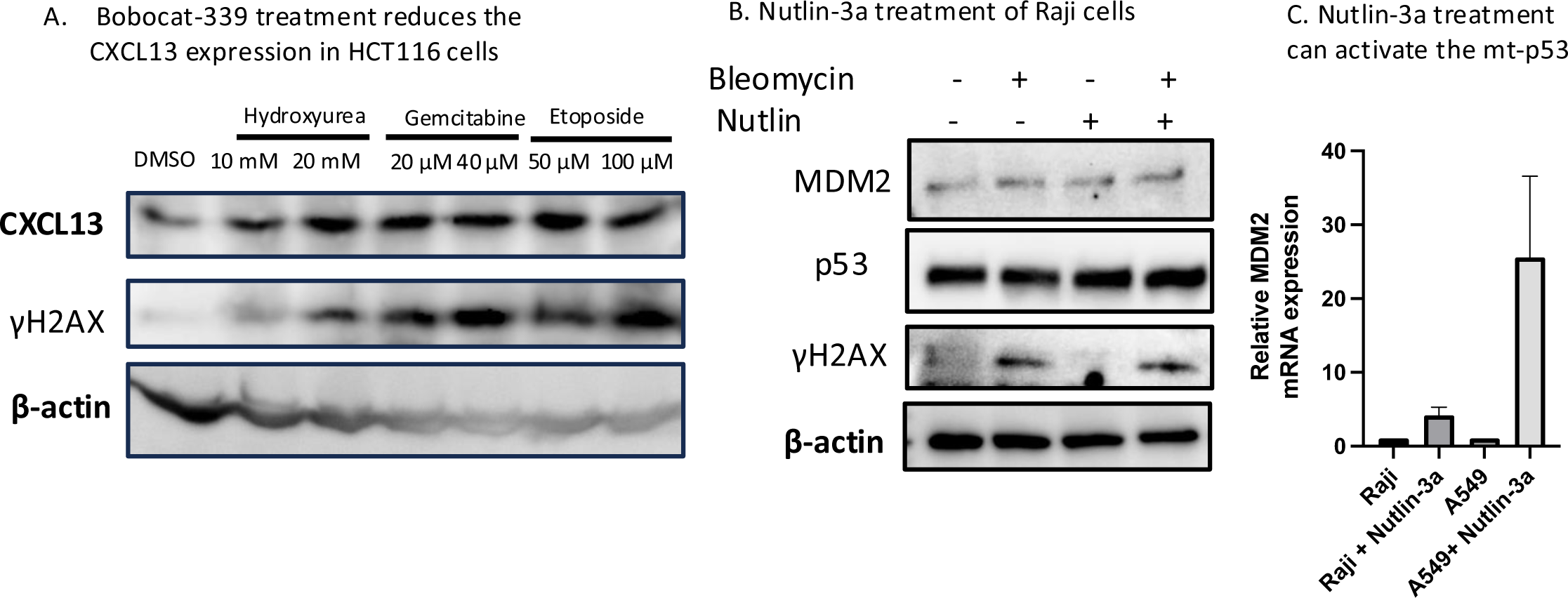
(A) Western blotting of CXCL13 and γH2AX from the cell lysates of HCT116 cells treated with DMSO, hydroxyurea (10 mM and 20 mM), Gemcitabine (20μM and 40μM) and Etoposide (50μM and 100μM) for 24 hours (B) Western blotting analysis of MDM2, p53, γH2AX and β-actin in Raji cells treated with Bleomycin (5μg/ml), Nutlin-3a (10μM), or the combination of Bleomycin (5μg/ml) + Nutlin-3a (10μM). (C) *MDM2* mRNA levels in the Raji/A549 cells treated with Nutlin-3a (10μM) for 24 hrs. Data are presented as mean ± SEM for n=3 samples.

**Supplementary Figure 3.**
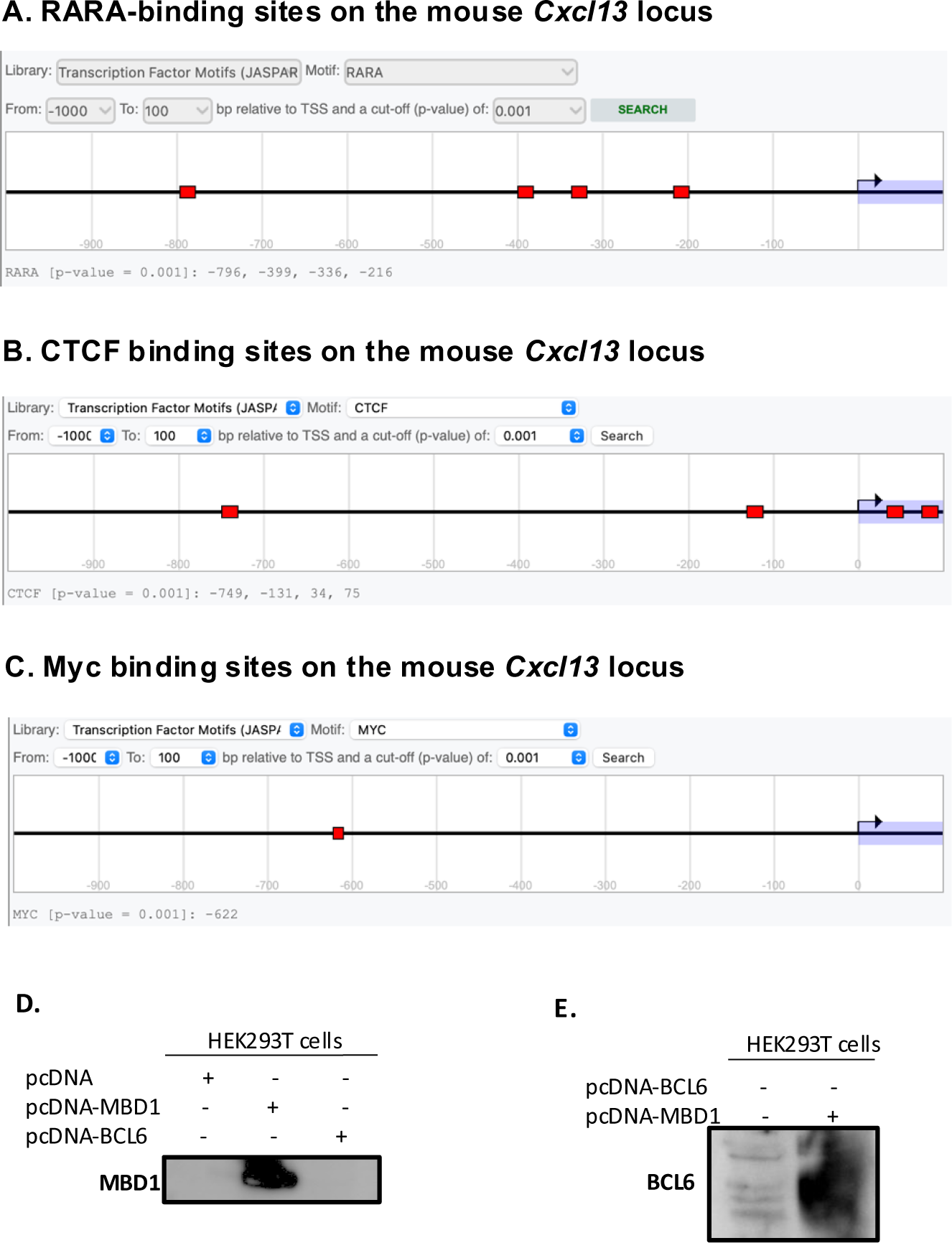
CTCF, RARA, Myc, and p53 binding sites are evolutionarily conserved on the *CXCL13* promoter (A) RARA binding sites are located at −216, −336, −399, and −796 from the Transcription Start Site (TSS) (B) Predicted CTCF binding sites are at 74, 34, −131, and −749 from the TSS (C) Predicted Myc binding site is at −622 from the *CXCL13*-TSS. (D) Western blotting of MBD1 in *pcDNA-MBD1* (1 μg) overexpressing H3K293T cells post 24 hr of transfection (E)Western blotting of BCL6 in *pcDNA-BCL6* (1 μg) overexpressing HEK293T cells post 24 hours of transfection.

**Supplementary Figure 4.**
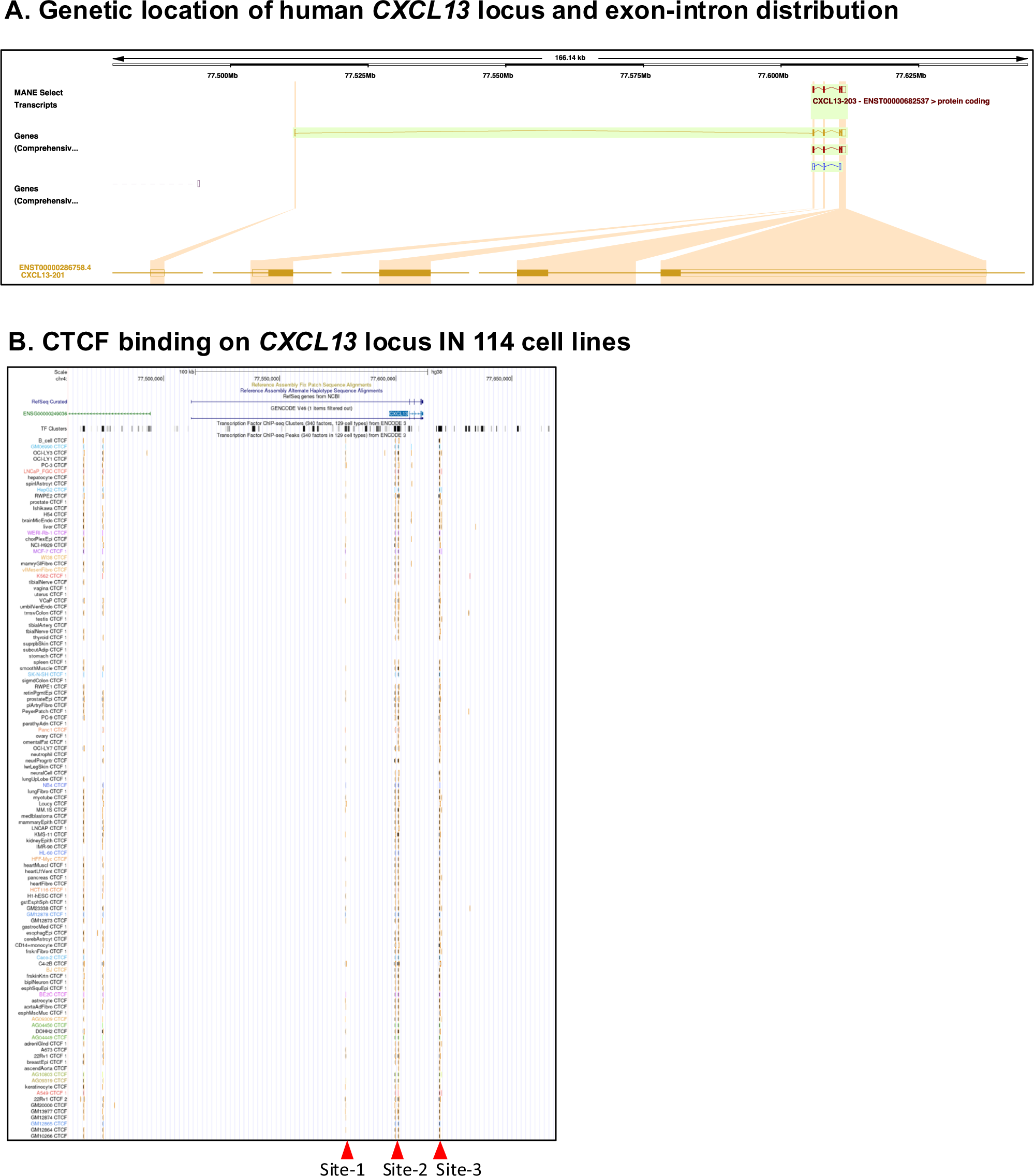
(A) CTCF-ChIP-seq and distribution of CTCF binding on site-1, site-2 and site-3 on CXCL13 locus in 114 cell lines including of normal human B cells and OCI-LY3. The additional peak of CTCF on site-3 is located at position, chr4:77606576-77606790. Peak 1 sequence Position: chr4:77578302-77578777. Position of peak 2 from: chr4:77599430-77599754 (B) Genomic Locus of human *CXCL13*. Ensemble Genome Browser view of the CXCL13 exon-intron position (transcript ID# ENST00000286758.4 CXCL13-201).

**Supplementary Figure 5.**
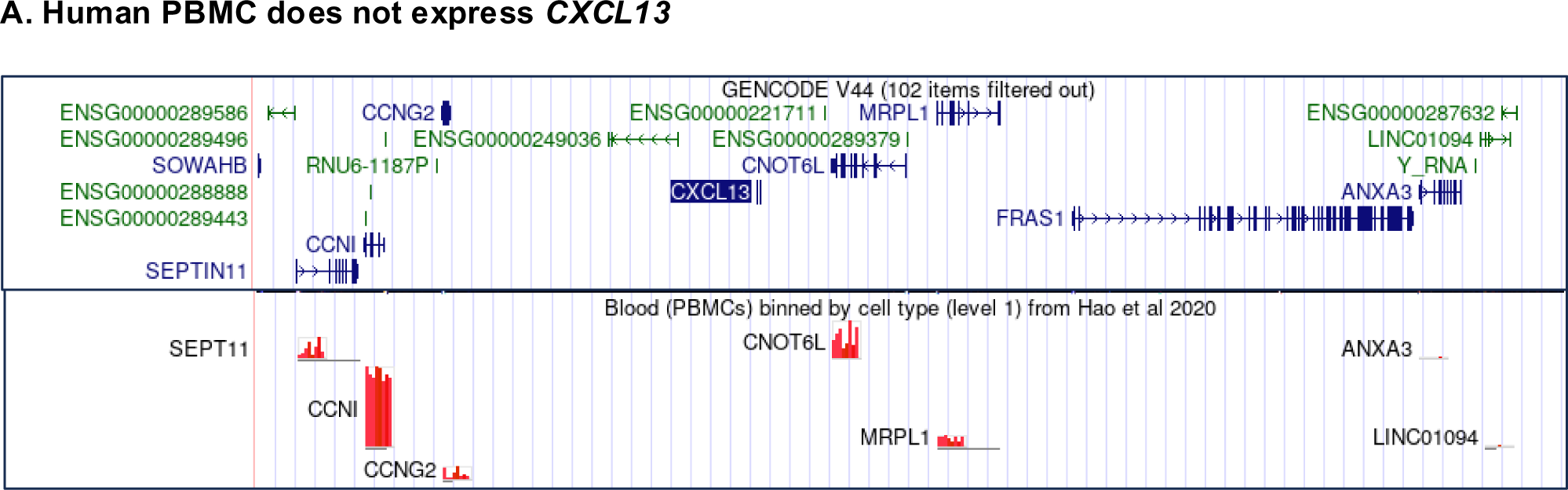
(A) Expression status of genes neighboring the *CXCL13* locus—SEP11, CCNI, CCNG2, CNOT6L, and MRPL1—in peripheral blood mononuclear B cells (PBMCs) is displayed, as assessed from PBMC data (Hao et al., 2020).

**Supplementary Figure 6.**
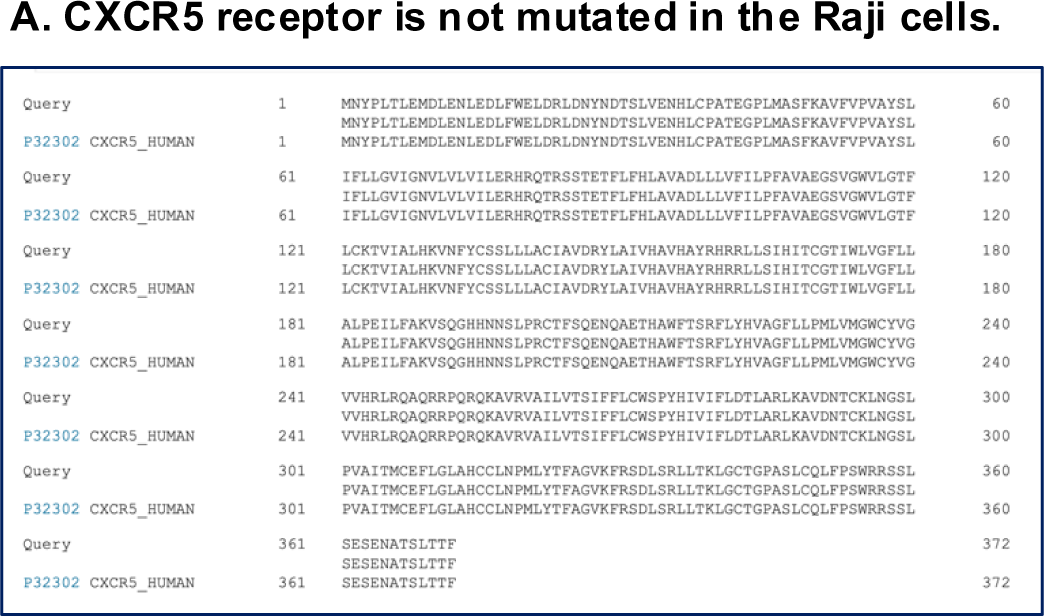
Sequencing of CXCR5 coding region in Raji cells (A) Absolute similarity in the peptide sequences derived from *CXCR5* DNA clones of Raji cells with human *CXCR5*. Uniport alignment of the CXCR5 protein from Raji cells and the human CXCR5 peptide from Uniprot. Three single nucleotide polymorphisms (SNPs) in Raji’s CXCR5 (Proline 39: CCC, Glycine 43: GGT, Threonine 338: ACC) did not resulted in amino acids confirming the reduced migration of Raji towards CXCL13 is not caused by *CXCR5* mutations.

**Supplementary Figure 7.**
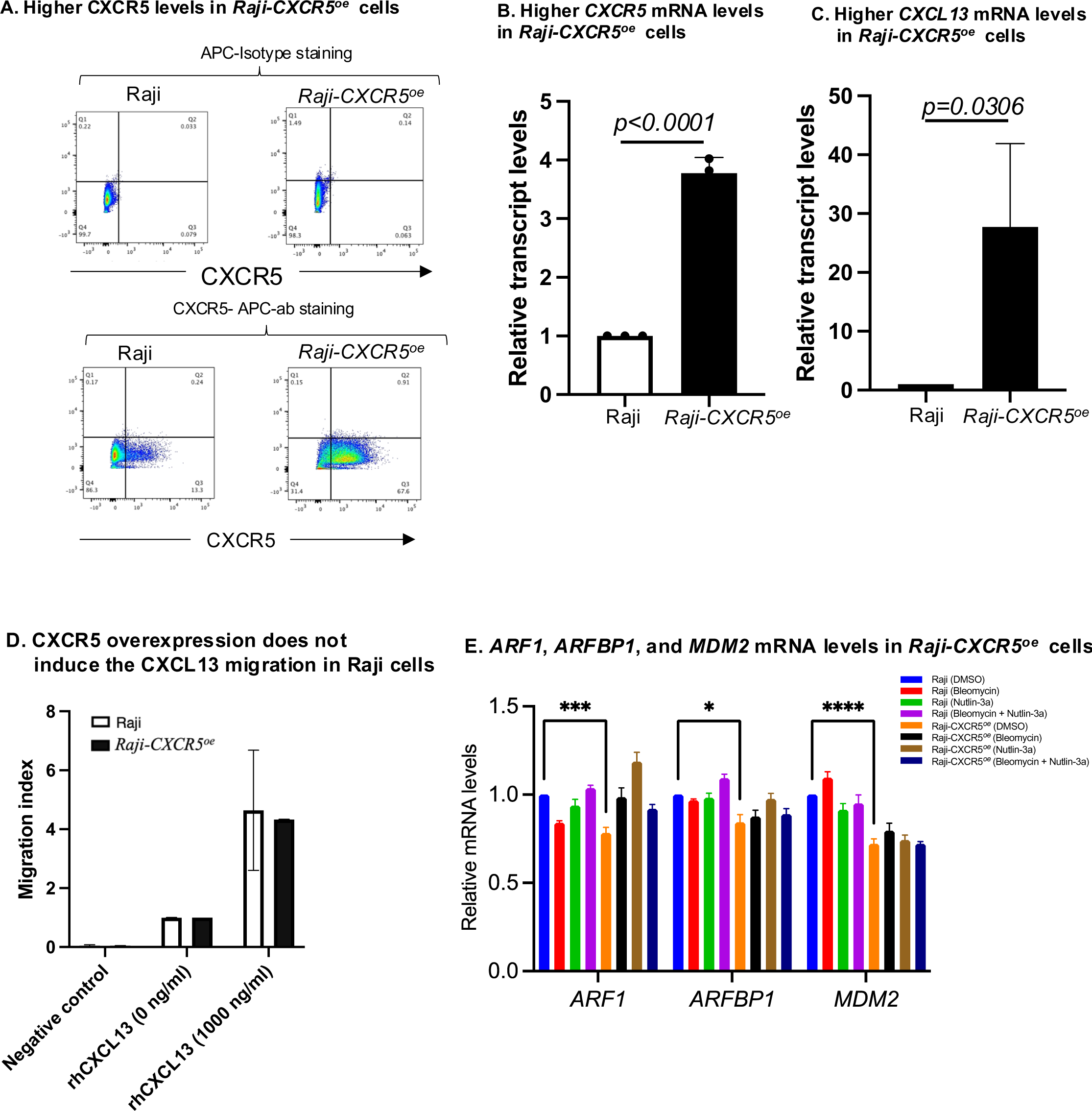
CXCR5 Overexpression induces *CXCL13* mRNA expression via positive feedback in Raji cells (A) Flow cytometry analysis of CXCR5 surface staining reveals elevated CXCR5 levels in *Raji-CXCR5^oe^* cells compared to normal Raji cells (B, C) Quantification of *CXCR5* and *CXCL13* transcripts using qRT-PCR in *Raji-CXCR5^oe^*cells compared to Raji cells. The data represent mean ± SEM for n=3, with p-values < 0.0001 and < 0.0306, respectively (D) Migration assays against rhCXCL13 gradient in Raji and *Raji-CXCR5^oe^* cells. Data presented as mean ± SEM for n=3 (E) qRT-PCR analysis of the expression levels of *ARF1*, *ARFBP1*, and *MDM2* in Raji and *Raji-CXCR5^oe^* cells after 24 hours treatment with either Bleomycin (5μg/ml), Nutlin-3a (10μM), or Bleomycin (5μg/ml) + Nutlin-3a (10μM). p= 0.0003 for Raji (DMSO) vs *Raji-CXCR5^oe^*, p= 0.0176 for Raji (DMSO) vs *Raji-CXCR5^oe^* and p<0.0001 Raji (DMSO) vs *Raji-CXCR5^oe^*. Tukey’s multiple comparison test. Data are mean ± SEM for n=3 samples.

**Supplementary table 1.**
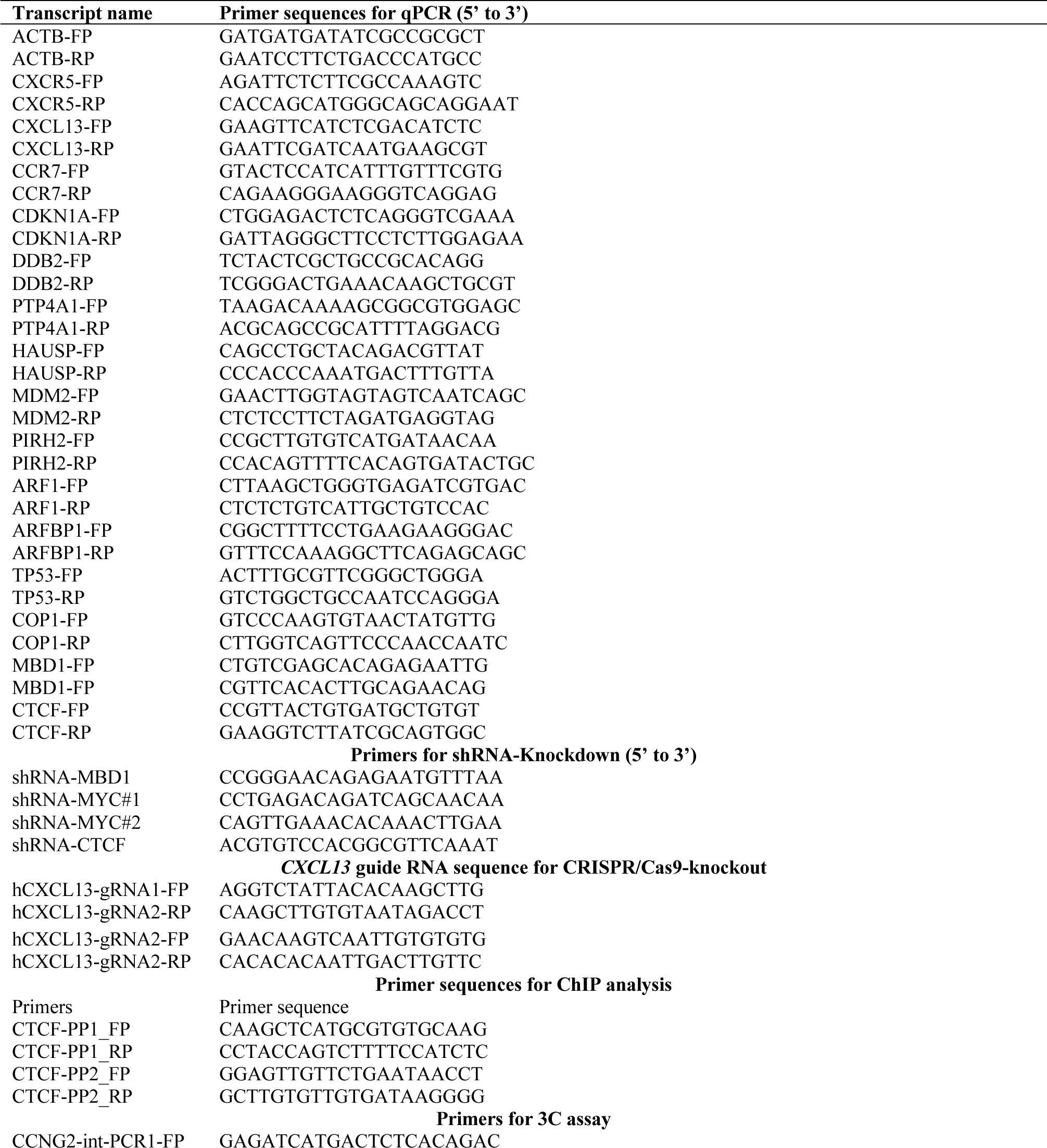
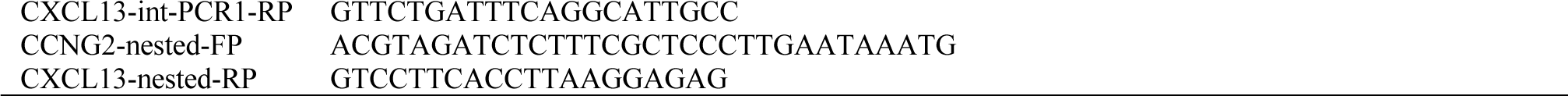
Primer sequences.

**Supplementary table 2.**
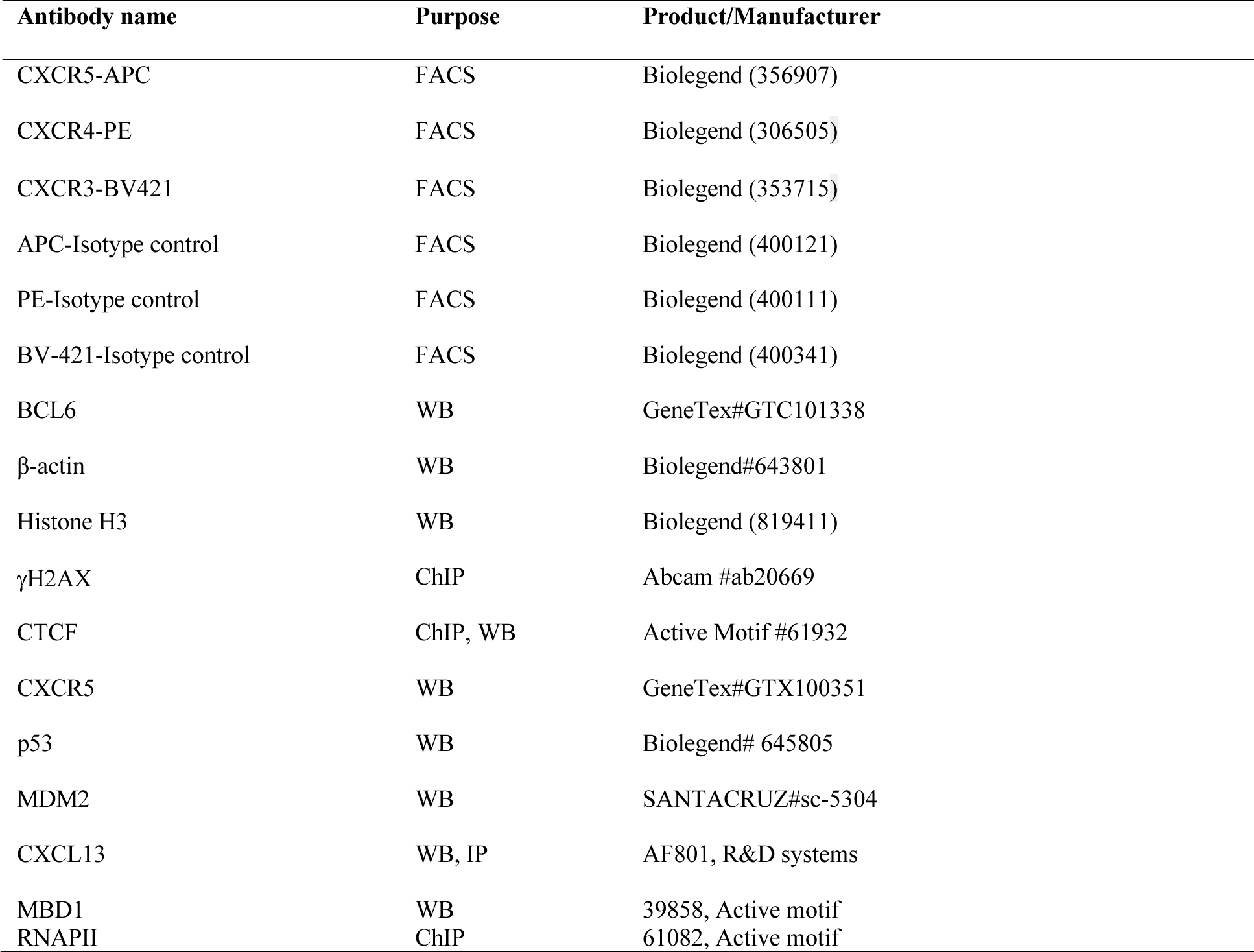
for antibodies for flow cytometry, western blotting, Immunoprecipitation and ChIP assays.

| Antibody name | Purpose | Product/Manufacturer |
| --- | --- | --- |
| CXCR5-APC | FACS | Biolegend (356907) |
| CXCR4-PE | FACS | Biolegend (306505) |
| CXCR3-BV421 | FACS | Biolegend (353715) |
| APC-Isotype control | FACS | Biolegend (400121) |
| PE-Isotype control | FACS | Biolegend (400111) |
| BV-421-Isotype control | FACS | Biolegend (400341) |
| BCL6 | WB | GeneTex#GTC101338 |
| β-actin | WB | Biolegend#643801 |
| Histone H3 | WB | Biolegend (819411) |
| γH2AX | ChIP | Abcam #ab20669 |
| CTCF | ChIP, WB | Active Motif #61932 |
| CXCR5 | WB | GeneTex#GTX100351 |
| p53 | WB | Biolegend# 645805 |
| MDM2 | WB | SANTACRUZ#sc-5304 |
| CXCL13 | WB, IP | AF801, R&D systems |
| MBD1 | WB | 39858, Active motif |
| RNAPII | ChIP | 61082, Active motif |

